# Genetic variation shapes human mRNA translation and disease risk

**DOI:** 10.64898/2026.02.10.705206

**Authors:** Siqi Wang, Chuyun Chen, Xinshu Xiao, Zefeng Wang

## Abstract

Genetic variation can influence protein abundance through translation, yet this regulatory layer remains poorly defined. We developed a deep learning approach to systematically map the effects of single-nucleotide variant (SNV) on translation efficiency (TE) across the human genome. In lymphoblastoid cells, >90,000 variants substantially altered TE, with strong positional and sequence-context biases. Importantly, the missense variants, traditionally considered only for their effect on protein recoding, were also found to reshape translation efficiency, with proline substitutions consistently reducing TE in a length-dependent manner. Extending our analysis to eight additional cell types revealed a two-layer architecture of translational regulation: 5’UTR variants produced highly concordant effects across cell types, whereas synonymous and missense variants in coding region exhibited cell type-specific outcomes, suggesting context-dependent translation regulation. TE-altering variants were enriched among GWAS loci and linked to cancer, immune, cardiometabolic, and neurological traits, positioning translation as a key mediator of genetic effects on disease.

## Introduction

Recent advances in high-throughput sequencing have facilitated comprehensive analyses of gene expression dynamics, uncovering the inconsistency between mRNA and protein abundances^1–4^ and highlighting the importance of translational regulation^5^. As the final step of gene expression, translation directly determines cellular protein abundance and allows cells to rapidly adapt to changing conditions, such as stress, apoptosis, and infection^6–8^. As such, translational control is essential to cellular function and holds significant implications for disease diagnosis, therapeutic development, and synthetic biology applications.

Translational regulation involves intricate interactions between *cis*-regulatory RNA elements and *trans*-acting factors that operate across both coding sequences (CDSs) and untranslated regions (UTRs), modulating the accessibility and recognition by the translational machinery^9–11^. In eukaryotes, this regulation occurs at multiple levels – initiation, elongation, and termination, with translation initiation typically being the rate-limiting step. Regulatory elements in the 5’UTR, such as sequence motif, upstream open reading frame (uORF), Internal Ribosome Entry Site (IRES), G-quadruplex (G4), and hairpin structure, are essential in modulating translation initiation^12–14^. In addition, features within the CDS and regulatory elements in the 3’UTRs also contribute to translational control and influence protein output^15,16^. These elements function synergistically or antagonistically, forming a dynamic and context-dependent regulatory network that ultimately determines translation efficiency. Dysregulation of translation is increasingly recognized as a key contributor to various human diseases such as cancer, infectious diseases, cardiovascular conditions, and neurodegenerative disorders^17–19^.

Given the central role of *cis*-regulatory elements in translation, genetic variants altering these elements may significantly affect translation outcomes and thus contribute to disease pathogenesis. Massively parallel reporter assays (MPRAs) have enabled functional testing of individual variants that affect translation^20,21^, but these assays rely on synthetic constructs and often fail to capture endogenous transcript contexts. In parallel, studies utilizing ribosome and polysome profiling have identified ribosomal occupancy quantitative trait loci (riboQTLs) or translation efficiency QTLs (teQTLs), and linked genetic variants to ribosome-related pathologies (ribosomopathies) and disease phenotypes^22–27^. However, QTL studies lack the resolution to pinpoint functionally causal variants. While early computational approaches have sought to predict functional variants affecting translation^28,29^, few methods offer a systematic evaluation of variant effects across full-length mRNAs. This gap is particularly notable given the vast number of potential regulatory variants and the limited scalability of experimental validation approaches.

Recent advances in deep learning have opened new opportunities for predicting the functional impact of genetic variants, especially in complex regulatory processes like translation. Inspired by natural language processing, deep learning architectures, such as convolutional neural network (CNN), recurrent neural network (RNN), and transformer, have been successfully applied to biological sequences to model local motifs, sequential dependencies, and interactions between *cis*-elements and *trans*-factors^30–36^. However, due to the high computational costs, most existing models are region-specific, focusing on the 5’UTR for translation initiation^37,38^, 3’UTR for RNA stability^39–41^, CDS for protein expression^42^, or short synthetic sequences tested by MPRAs^43^. While these models have yielded valuable insights, they often overlook the broader contextual and combinatorial regulation that occurs across full-length transcripts.

To address these challenges, we developed TEFL-mRNA, a hybrid deep-learning model integrating CNN and RNN architectures to predict <u>T</u>ranslation <u>E</u>fficiency (TE) from Full-<u>L</u>ength <u>mRNA</u> sequences. We applied TEFL-mRNA to ribosome profiling data across nine human cell types and systematically assessed the translational impact of millions of single-nucleotide variants. This analysis revealed tens of thousands of variants with significant TE-altering effects, uncovered both shared and cell type-specific patterns of variant effects, and highlighted strong enrichment of functional variants in immune and cancer pathways. Together, our study establishes translation as a widespread target of genetic variation and provides a generalizable framework for variant interpretation in human disease.

## Results

TEFL-mRNA model for predicting translation efficiency from full-length mRNA sequences

To develop a model for mRNA translation, we first constructed a training set by calculating TE values of individual transcripts using ribosome profiling and RNA-seq datasets from 47 human lymphoblastoid cell lines (LCLs). After applying a series of preprocessing filters, including removing transcripts with a high proportion of missing values or low read counts, as well as discarding samples with low pairwise correlations, we classified the transcripts into three categories with high, low, and intermediate TE (see Methods) (**Figure 1A**). Next, we embedded full-length mRNA sequences using one-hot encoding, with an additional row to label mRNA regions (‘0’ for 5’UTR, ‘1’ for CDS, and ‘2’ for 3’UTR). The resulting matrix was used as an input to a hybrid neural network composed of a CNN and an RNN, followed by a dense layer (**Figure 1B**, Supplementary Figure 1). The entire model, collectively termed TEFL-mRNA, was then trained to predict TE categories.

**Figure 1.**
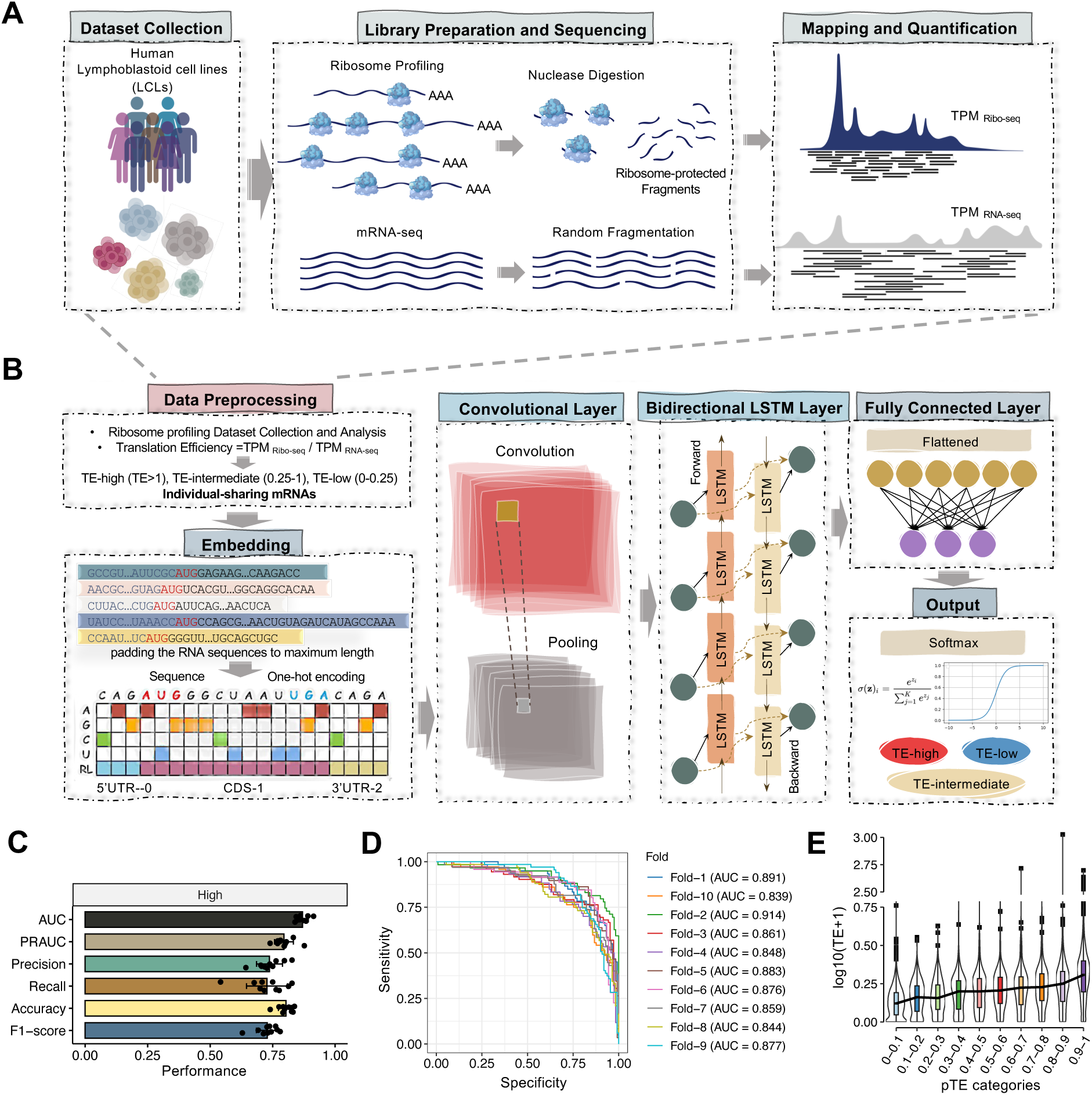
Construction and evaluation of the TEFL-mRNA model. (**A**) Overview of ribosome profiling and RNA-seq data processing. Transcripts per million (TPM) values were calculated from Ribo-seq and RNA-seq data, and translation efficiency (TE) was calculated as TPM_ribo-seq_/TPM_RNA-seq_. mRNAs were classified into high-, intermediate-, and low-TE categories, with “individual-sharing” transcripts defined by consistent classification across samples after filtering (see Methods). (**B**) Architecture of the convolutional-recurrent hybrid neural network for TE prediction. Full-length mRNA sequences were one-hot encoded, with a fifth channel labeling RNA regions (5’UTR, CDS, and 3’UTR), and fed into the CNN and BiLSTM layers, followed by fully connected layers to output the predicted TE-high probabilities (referred to as pTE). (**C**) Model performance metrics (AUC, PRAUC, Precision, Recall, Accuracy, and F1-score) from 10-fold cross-validation; each point represents one-fold. (**D**) Receiver operating characteristic (ROC) curves for the 10 folds; the x-axis shows specificity; the y-axis shows sensitivity. (**E**) Relationship between predicted pTE categories and actual TE values. The x-axis shows predicted score categories; the y-axis shows log_10_(TE+1) across samples to accommodate zero values.

Although the multi-class training may capture a richer spectrum of TE variation that includes more nuanced features, our primary interest lies in the analysis of highly translated transcripts, which are often critical drivers of cellular protein synthesis and are more useful in design of mRNA drugs. Therefore, we evaluated the model’s ability to accurately identify highly translated mRNA (TE-high *vs.* others). Using the predicted TE-high probability (referred to as pTE hereafter), we calculated several performance metrics, including AUC, PRAUC, precision, recall, accuracy, and F1 score. Overall, this model achieved an AUC above 85% (average AUC = 0.869) and an accuracy of 80% (average accuracy = 0.803) across 10-fold cross-validation (Figures 1C, D), suggesting the TEFL-mRNA model can reliably identify the transcripts with high TE. In addition, the pTE values showed a positive correlation with experimentally measured TE (Figure 1E). Based on this relationship, we use the pTE as a proxy for TE values in the downstream analyses of translational regulation.

### TEFL-mRNA recapitulates known mechanisms of translational regulation

To assess whether TEFL-mRNA captures known biological mechanisms, we tested its ability to recognize canonical *cis*-regulatory features involved in translational inhibition. miRNAs are well known to mediate gene silencing by promoting mRNA degradation or inhibiting translation, typically through binding to the 3’UTR^44^. Consistent with this regulatory mode, we found that transcripts harboring miRNA binding sites in their 3’UTRs had significantly lower pTE scores compared to those without binding sites (Figure 2A). Interestingly, transcripts with miRNA binding sites in the 5’UTR displayed higher pTE scores than those lacking 5’UTR binding sites (Supplementary Figure 2A, right panel), suggesting a potential context-specific role for miRNAs in enhancing translation in agreement with previous literature^45–47^.

**Figure 2.**
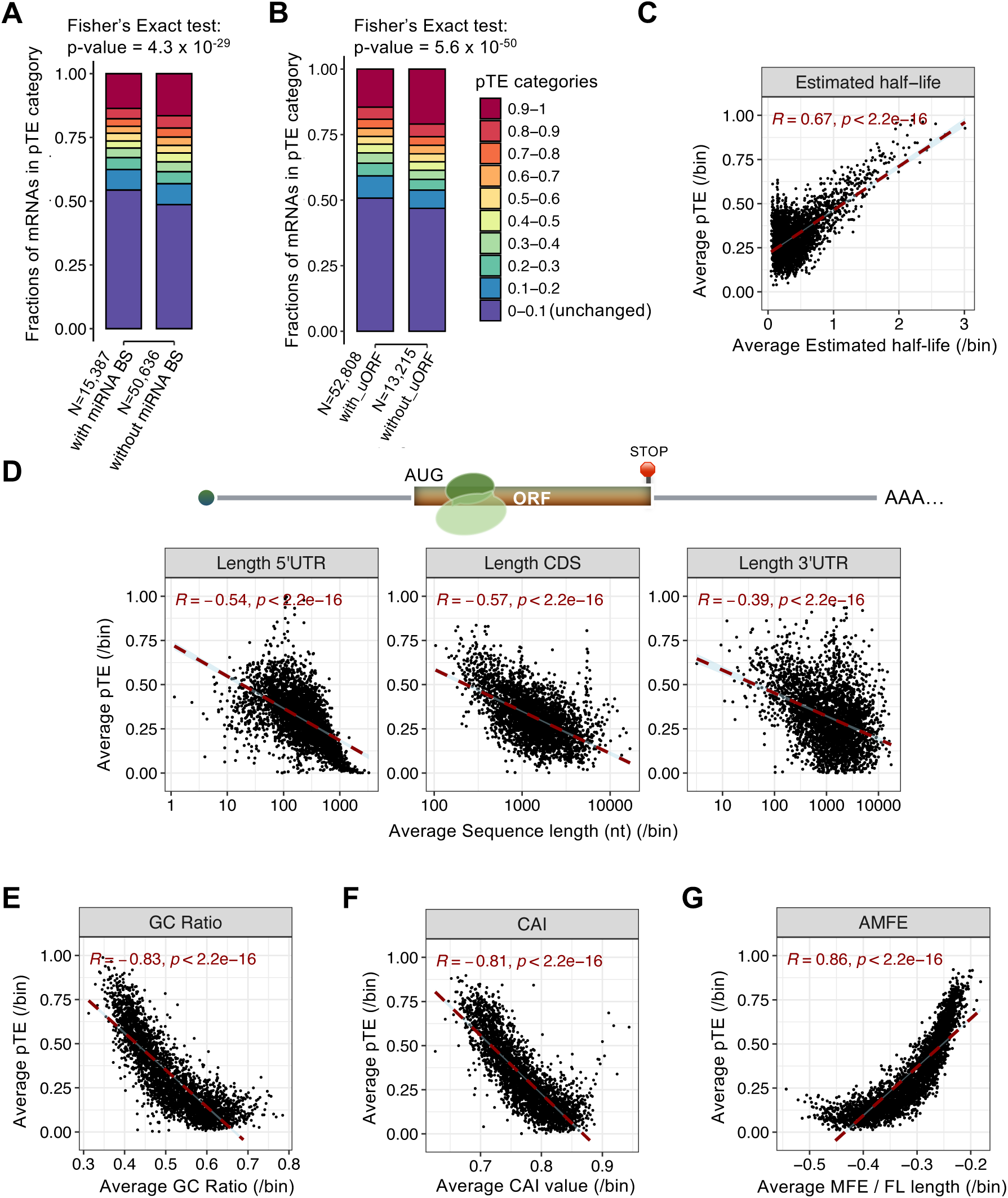
**Sequence and structural correlates of translation efficiency captured by TEFL-mRNA.** (**A-B**) Distribution of pTE across mRNAs with or without (A) 3’UTR miRNA binding sites or (B) 5’UTR upstream open reading frame (uORF). Y-axis shows the fraction of mRNAs in each pTE category. P-value from Fisher’s Exact test indicates significant shifts in pTE distributions. (**C**) Correlation between estimated mRNA half-life and predicted TE. (**D**) Correlations between pTE and the length of the 5’UTR, CDS, and 3’UTR. (**E-G**) Correlations between pTE and (E) GC ratio, (F) codon adaptation index (CAI), or (G) adjusted minimum free energy (AMFE; minimum free energy normalized by mRNA length). For all scatter plots, points represent bin-averaged (10 mRNAs per bin) values of sequence features (x-axis) and pTE (y-axis); Pearson’s correlation coefficients (R) and p values are shown.

uORFs in the 5’UTR are well-established regulators of translation, typically repressing translation of the main open reading frame (mORF)^48,49^. To assess whether our model captures this regulatory mechanism, we analyzed the transcripts harboring uORFs, and found that such transcripts were significantly enriched among those with pTE < 0.5 (**Figure 2B**) and showed lower overall pTE scores (Supplementary Figure 2B). These findings are consistent with the known repressive roles of uORFs and further support the model’s ability to capture established mechanisms of translational regulation.

### TEFL-mRNA reveals global sequence and structural correlates of translation

Having shown that TEFL-mRNA captures canonical regulatory mechanisms, we next examined whether the model could identify broader sequence and structural features associated with translation efficiency. We examined a range of transcript-level properties, including mRNA half-life, lengths of mRNA regions, codon usage (measured by the Codon Adaptation Index, CAI), GC ratio, and mRNA structures (estimated by Adjusted Minimal Free Energy, AMFE) (see Methods).

We found that transcripts with high pTE values are significantly correlated with longer half-lives (**Figure 2C**), consistent with previous reports of translation-dependent mRNA degradation^50,51^. In contrast, longer mRNAs generally exhibited lower pTE (**Figure 2D**, Supplementary Figure 3A), likely due to the increased presence of regulatory elements, such as uORFs and miRNA binding sites distributed throughout the mRNA regions, as well as a greater propensity to form secondary or tertiary structures that impede translation^52–54^.

Previous studies have linked GC content, mRNA secondary structure, and codon optimality positively with translation efficiency, primarily through their effects on mRNA stability and tRNA availability^55^. Surprisingly, we observed that both GC content and CAI were negatively correlated with pTE, whereas AMFE showed a positive correlation (**Figure 2E-G**, Supplementary Figure 3B-D). To confirm that our observations were not TE-prediction artifacts, we repeated the analyses using experimentally measured TE values from ribosome profiling and observed a consistent trend (Supplementary Figure 4). These results suggest that in human LCLs, the relationship between sequence features and translation efficiency may differ from canonical expectations, pointing to potential context-specific constraints on codon usage and RNA structure (see Discussion).

### TEFL-mRNA identifies genetic variants that alter translation at single-nucleotide resolution

After characterizing global sequence and structural correlates of translation, we next applied TEFL-mRNA to examine how individual sequence variant may influence translation efficiency. For each SNV, we assessed its pTE difference between the reference and alternative alleles (pTE_alt_ – pTE_ref_, or ΔpTE) (**Figure 3A**). To define a threshold for the substantial TE alteration, we used a control set of common non-GWAS, non-ClinVar dbSNP variants (allele frequency 0.1-0.9) as a proxy for putatively neutral variation. We examined the distribution of |ΔpTE| values in this set and selected a cutoff of 0.1, corresponding to 99.73% (i.e., 0.27% tail probability) of the control distribution (Supplementary Figure 5A). This definition provides a conservative threshold expected to yield a false positive rate of 0.27% when applied to neutral variants. We then applied this threshold to all annotated 5’UTR, 3’UTR, synonymous, and missense SNVs in the gnomAD database, identifying 90,644 SNVs as putative TE-altering variants (Supplementary Table 1).

**Figure 3.**
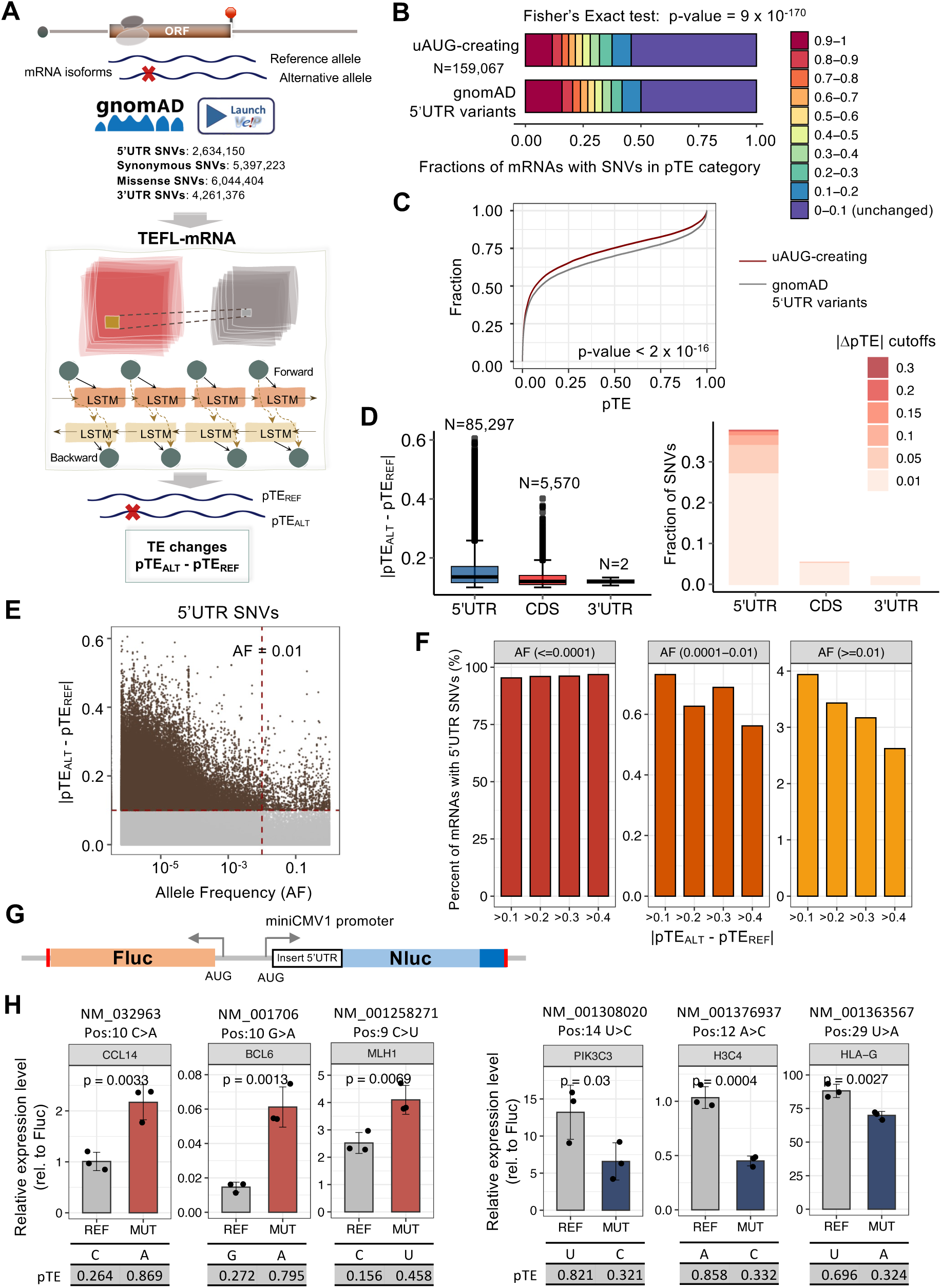
**The effects of single-nucleotide variants on translation efficiency.** (**A**) Overview of TEFL-mRNA variant analysis. Genetic variants from gnomAD were annotated with VEP, and mRNA transcripts containing 5’UTR, 3’UTR, or CDS were extracted. Full-length mRNA sequences with reference and alternative alleles were input into TEFL-mRNA to calculate TE changes (ΔpTE = pTE_ALT_ – pTE_REF_). (**B**) Distribution of pTE categories for mRNAs with uAUG-creating SNVs versus all gnomAD 5’UTR variants; enrichment tested by Fisher’s Exact test. X-axis shows the fraction of mRNAs in each pTE category. (**C**) Cumulative distribution of pTE for uAUG-creating (red line) versus gnomAD 5’UTR (grey line) variants; p-value from Kolmogorov-Smirnov test. (**D**) Regional distribution of TE effects for TE-altering variants. Left: boxplot of |ΔpTE| of variants with |ΔpTE| > 0.1 in different regions. The number of such variants is shown above each box. Right: fraction of SNVs passing different |ΔpTE| cutoffs. (**E**) Relationship between allele frequency (AF) and |ΔpTE| for 5’UTR SNVs. Brown points mark variants with |pTE| > 0.1. (**F**) Percentage of transcripts with 5’UTR SNVs passing the specified |ΔpTE| cutoff within the allele frequency (AF) range among all transcripts with 5’UTR SNVs that pass the |ΔpTE| cutoff. (**G**) Schematic of the Fluc-Nluc dual luciferase reporter used to assay SNV effects on TE. (**H**) Reporter assay results for six gnomAD SNVs. Left: SNVs whose variant alleles increased TE; right: SNVs whose variant alleles decreased TE. Bars show mean ± standard deviation (SD) from biological triplicates. Y-axis shows the normalized luciferase ratio. P values from one-tailed t-tests are shown. TEFL-mRNA predicted pTE values are shown below each panel.

As a validation, we first focused on variants predicted to create uAUGs in the 5’UTR, which are known to introduce alternative translation initiation sites and suppress translation from the canonical start codon^56^. As expected, TEFL-mRNA predicted significantly lower pTE scores for the mRNA transcripts that carry uAUG-creating variants compared to all 5’UTR SNVs (**Figure 3B**, C). These results support the model’s ability to capture biologically relevant regulatory effects of genetic variants on translation. Notably, a large number (and higher fraction) of SNVs in the 5’UTRs were predicted to alter TE, exhibiting greater |ΔpTE| values than variants in other regions (**Figure 3D**). Furthermore, among the predicted functional 5’UTR SNVs, we observed a general trend of higher |ΔpTE| for variants with lower allele frequency (**Figure 3E**), with the strongest effects seen in SNVs with extremely low allele frequencies (<0.0001). When applying increasingly stringent |ΔpTE| thresholds, rare variants were disproportionately retained, while the more common variants progressively dropped out (**Figure 3F**, Supplementary Figure 5B), suggesting that rare SNVs are more likely to drive substantial changes in translation.

To experimentally validate TEFL-mRNA predictions, we designed a dual luciferase reporter system containing an Rluc driven by a 5’UTR carrying either the reference or alternative allele of an SNV, and a Fluc gene as an internal control (Figure 3G). The resulting constructs were transfected into cultured cells to quantify the relative changes of translation by measuring luminescence intensities (see Methods). Consistent with model predictions, the tested SNVs significantly altered luciferase activity. The alternative alleles in the tested genes *CCL14*, *BCL6*, and *MLH1* enhanced translation (Figure 3H, left panel), while those in *PIK3C3*, *H3C4*, and *HLA-G* decreased TE (Figure 3H, right panel). These genes span diverse functional categories, including immune signaling, autophagy, chromatin regulation, and DNA repair, many of which are known to intersect with pathways dysregulated in diseases such as cancer^57–62^. These findings not only support the predictive accuracy of TEFL-mRNA, but also provide a foundation for the broader disease-relevant analyses described below.

### TE-altering SNVs showed sequence and positional biases across mRNAs

Having established that a large number of SNVs can impact TE, we next examined whether such functional SNVs exhibited different distribution by alleles and positions in the mRNA. We focused on variants within ±1 kb of the start and stop codons that are known to contain regulatory elements. Our analysis revealed clear region- and allele-dependent translation effects of SNVs (**Figure 4A**). First, the variants in the 5’UTR exerted the strongest impact, particularly those near the start codon, highlighting the importance of initiation in translation control. Second, the allele identity also influenced TE changes. Variants introducing A or U alleles were generally associated with increased pTE, whereas C or G alleles tended to reduce it. This trend was especially pronounced in the 5’UTR, where more stringent ΔpTE thresholds further accentuated the enrichment of A alleles among TE-increasing variants and C/G alleles among TE-decreasing ones (**Figure 4B**). This observation is consistent with our earlier finding that higher GC content correlated with lower pTE (**Figure 2E**).

**Figure 4.**
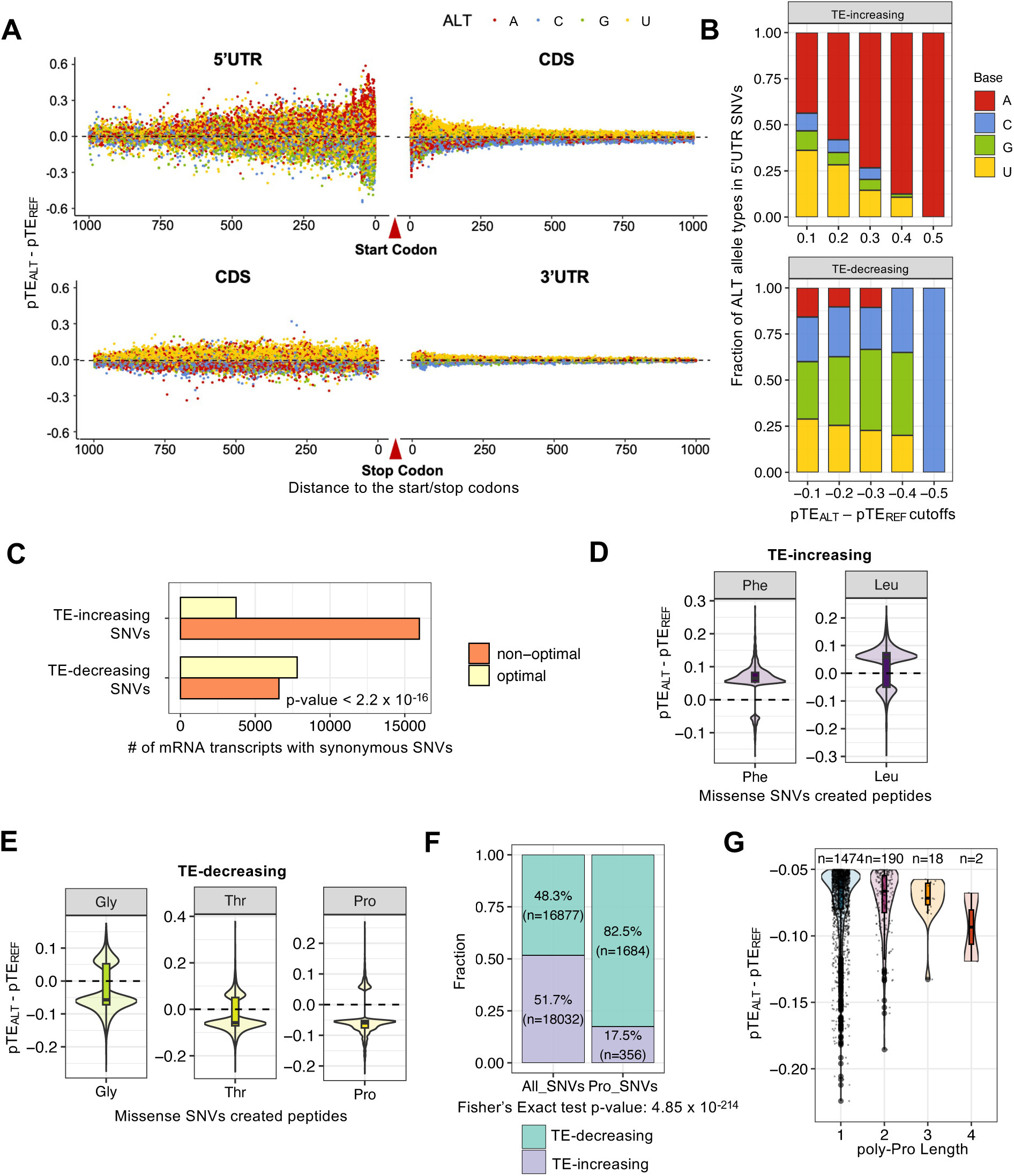
**TE-altering SNVs in coding and noncoding regions.** (**A**) Predicted ΔpTE for SNVs located within ±1000nt of the start or stop codon. Each point indicates an individual SNV; colors indicate the alternative allele. X-axis shows the relative distance to the start or stop codon. (**B**) Fractions of transcripts with 4 alternative allele types (A/C/G/U) among 5’UTR SNVs exceeding different ΔpTE cutoffs. (**C**) Counts of TE-increasing and TE-decreasing synonymous SNVs with |ΔpTE| > 0.05 (corresponding to a false positive rate of 1%, similarly in panels D-G below). Variants changing codons toward non-optimal (orange) or optimal (yellow) are shown separately. P-value from Fisher’s exact test. (**D-E**) ΔpTE for missense SNVs introducing specific amino acids. (D) TE-increasing variants frequently introduce hydrophobic residues (Phe, Leu). (E) TE-decreasing variants more often introduce Gly, Thr, and Pro, linked to reduced elongation efficiency. (**F**) Proportion of TE-increasing versus TE-decreasing effects among all missense SNVs or Pro-substituting SNVs; enrichment tested by Fisher’s Exact test. Y-axis shows the fraction of transcripts with the SNVs in each category. (**G**) Relationship between ΔpTE and poly-proline tract length created by missense SNVs. n denotes the number of SNVs per group.

### Synonymous and missense variants have TE-altering potential

We next examined synonymous SNVs, which preserve the amino acid sequence but can influence translation through codon usage. To increase the sensitivity of our analysis and capture a broader set of codon changes, we used a relaxed threshold |ΔpTE| > 0.05 (i.e., false positive rate of 1%, Supplementary Figure 5A) for the CDS SNV analysis. Substitutions toward codons deemed ‘optimal’ by CAI did not uniformly enhance TE; many were associated with reduced TE (**Figure 4C**), with effects varying by amino acid and codon context. Synonymous variants that decreased GC content in codons were more likely to increase TE (Supplementary Figure 6A, B). These codon-level patterns are consistent with our gene-level analysis showing a negative correlation between CAI or GC content and TE in LCLs (**Figure 2E**, F), indicating that higher CAI does not necessarily imply higher translation efficiency in this context (see Discussion).

Although missense variants have traditionally been studied for their impact on protein sequence and function, our analysis reveals that they can also exert substantial effects on translation. We identified 22,532 missense variants with |ΔpTE| > 0.05 and observed clear amino acid-specific biases (Supplementary Figure 6C). Specifically, SNVs introducing leucine (Leu) and phenylalanine (Phe) were predominantly associated with increased TE (**Figure 4D**), whereas TE-decreasing SNVs more often introduce threonine (Thr), glycine (Gly), and proline (Pro) (**Figure 4E**). In particular, proline substitutions showed a strong and consistent inhibitory effect: 82.5% of TE-altering SNVs introducing proline reduced TE, a highly significant enrichment compared to all TE-altering missense SNVs (Fisher’s exact test p < 4.85 × 10⁻²¹⁴, **Figure 4F**). This inhibitory effect intensified with increasing poly-proline tract length in the peptide region surrounding the variant, with progressively lower pTE observed for longer runs of prolines (**Figure 4G**). This observation supports previous findings that consecutive proline residues stall ribosomes due to their conformational rigidity, which impedes peptide bond formation and slows translation^63,64^.

### TE-altering SNVs may modulate translation via mRNA structural remodeling

We next examined whether TE-altering SNVs impact local RNA secondary structure, which plays a key role in ribosome loading and scanning^13^. Because predicting RNA folding across millions of sequences is computationally intensive, we focused on a well-curated subset of SNVs from the ExAC database, which includes 60,706 human exomes. From this dataset, we identified TE-altering SNVs in the 5’UTR and CDS regions as those with |ΔpTE| > 0.1, and defined TE-unchanged control SNVs as those with negligible |ΔpTE| (< 10^-6^). RNA structure shifts were then predicted for each SNV by comparing secondary structures between the alternative and reference alleles. Notably, both TE-increasing (i.e., ΔpTE > 0.1) and TE-decreasing (i.e., ΔpTE < −0.1) SNVs induced significantly larger structural shifts than TE-unchanged SNVs (**Figure 5A**, Supplementary Table 2).

**Figure 5.**
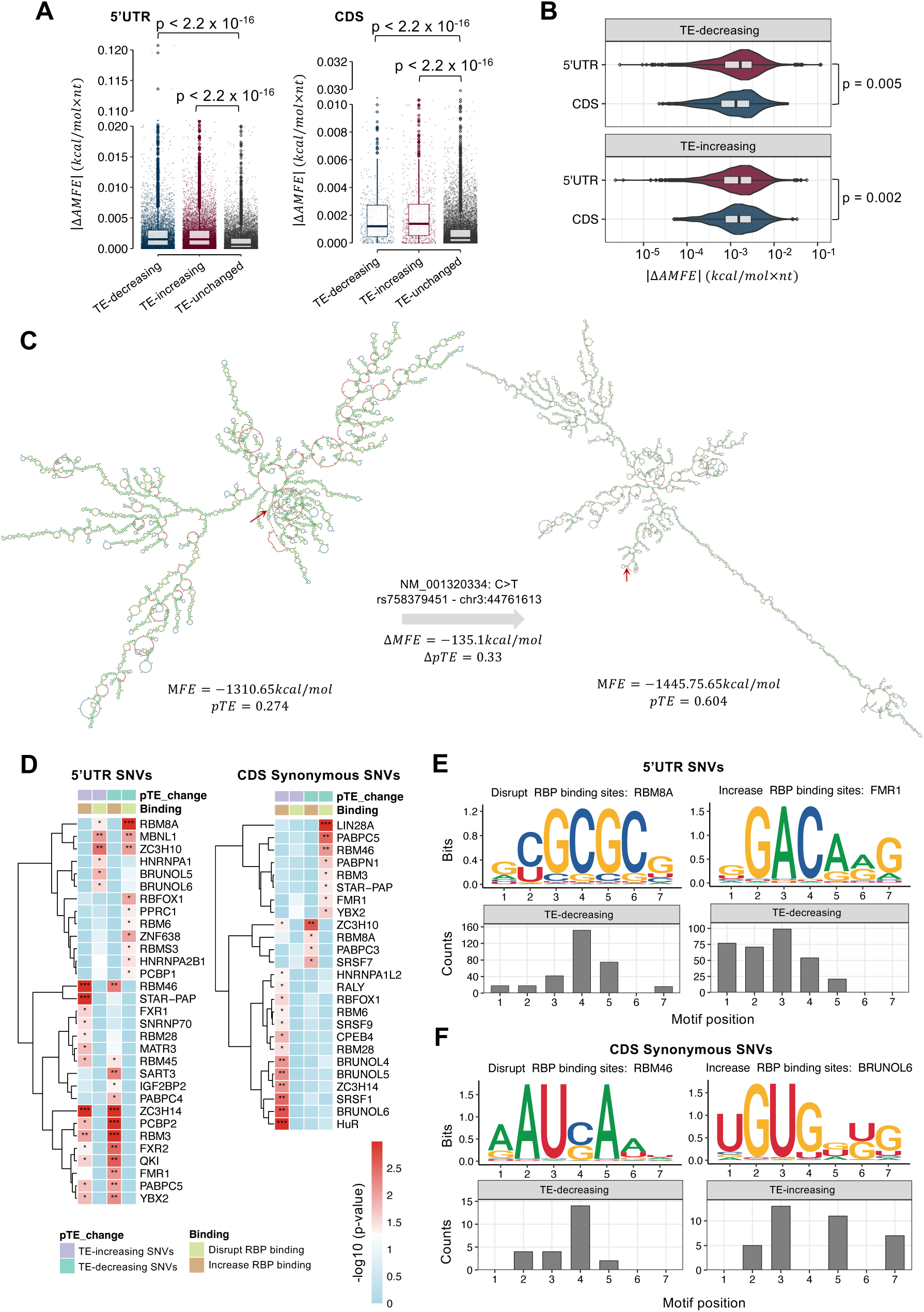
**Contribution of mRNA structure and RBP binding to translation alteration.** (**A**) Absolute changes in adjusted minimum free energy (|ΔAMFE|) for TE-decreasing, TE-increasing, and TE-unchanged SNVs in the 5’UTR and CDS. (**B**) Comparison of |ΔAMFE| between TE-decreasing and TE-increasing SNVs in the 5’UTR and CDS; p-values from Student’s t-test are shown. (**C**) Structures of the mRNA NM_001320334 with the reference (C, left) or alternative allele (T, right) at an SNV (rs758379451) in the 5’ UTR. Structures were predicted using LinearFold and visualized with foRNA. (**D**) Heatmaps showing RBPs whose binding was significantly changed by TE-altering SNVs (compared to TE-unchanged SNVs) in 5’UTR (left) and CDS (right). Binding changes were predicted by DeepBind; p-values from Student’s t-test are color-coded. (**E-F**) Representative RBP binding motifs in the 5’UTR (E) and CDS (F). For each motif, bar plots show counts of TE-altering SNVs that map to specific motif positions.

Furthermore, TE-altering variants in the 5’UTR induced larger structural changes than their CDS counterparts (Figure 5B), consistent with the pivotal role of the 5’UTR in translation initiation. One illustrative example is shown in Figure 5C, where a 5’UTR SNV in the *KIAA1143* transcript was predicted to increase TE and stabilize the mRNA structure. *KIAA1143* encodes a protein associated with cell growth and motility^65^, highlighting the potential biological impact of structural modulation. These findings emphasize the role of RNA secondary structure as a key regulatory layer in translational control, particularly within the 5’UTR, which suggests that some TE-altering SNVs may exert their effects by reshaping mRNA folding landscapes.

### TE-altering SNVs may impact translation through altered protein-RNA binding

Given the central role of RNA-binding proteins (RBPs) in translational control^10^, we next examined whether TE-altering SNVs may modulate translation by altering RBP binding. Using DeepBind^66^, a deep learning model that predicts RBP binding affinity, we compared the binding scores of reference and alternative alleles for each SNV (see Methods). TE-altering SNVs in the 5’UTR and CDS showed significant changes in the predicted RBP binding compared to TE-unchanged SNVs (**Figure 5D**). A subset of these SNVs demonstrated dual effects, enhancing the binding of certain RBPs while disrupting the interaction with others (Supplementary Figure 7, Supplementary Table 3).

Depending on the function of the RBP, altered binding could either repress or enhance translation. For example, RBM8A, a core protein in the exon junction complex (EJC) linked to mTOR signaling and nonsense-mediated decay^67^, showed reduced binding at SNVs in the 5’UTR, which was associated with decreased TE (Figure 5D, left panel). Similarly, increased binding of FMRP (encoded by *FMR1*), a known translation repressor^68^, was also linked to reduced TE. These effects were position-dependent: ‘C’ at the 4th position of RBM8A binding motif (GCGCGCG), and ‘A’ at the 3rd position of FMRP binding motif (GGACAAG), were often affected by TE-altering SNVs (Figure 5E). In general, both weakened and strengthened binding of some RBPs could enhance or repress TE depending on the gene target (Figure 5D), reflecting context-dependent RBP function.

Similarly, in the CDS, we observed both disrupted and enhanced RBP binding (**Figure 5D**, right panel). SNVs that disrupted binding of RBPs with A-rich motifs, including RBM46 (AAUCAAU), PABPC5 (AGAAAAU), and LIN28A (GAAGGAA), tended to reduce TE. In contrast, variants that increased binding of UG-rich RBPs such as the BRUNOL family (UGUGGUG/UGUGUGU) and SRSF1 (GGAGGA) were associated with enhanced TE. Many of these effects centered on core positions of the motifs, such as the third ‘U’ in the BRUNOL6 motif (**Figure 5F**). Interestingly, the same RBP could have opposing effects on TE depending on whether binding occurred in the 5’UTR or CDS, again reflecting context-dependent regulation of translation by RBPs. Together, these findings reveal that single-nucleotide changes may fine-tune TE by modulating RBP binding in a region- and context-specific manner.

### TE-altering SNVs are associated with human diseases and GWAS traits

To assess the functional significance of TE-altering SNVs, we conducted gene and disease ontology enrichment analyses focusing on SNVs with |ΔpTE| > 0.1. This analysis revealed strong enrichment in cancer-related biological processes, including the G2/M cell cycle transition, intrinsic apoptotic signaling, and response to oxidative stress. Notably, we also observed enrichment in antigen processing and presentation via MHC class I, a key pathway in immune defense and cancer immunology (**Figure 6A**). Disease ontology analysis further highlighted the relevance of TE-altering SNVs to multiple cancer types (**Figure 6B**), supporting the contribution of translational regulation to disease mechanisms, particularly in oncogenesis and immune function.

**Figure 6.**
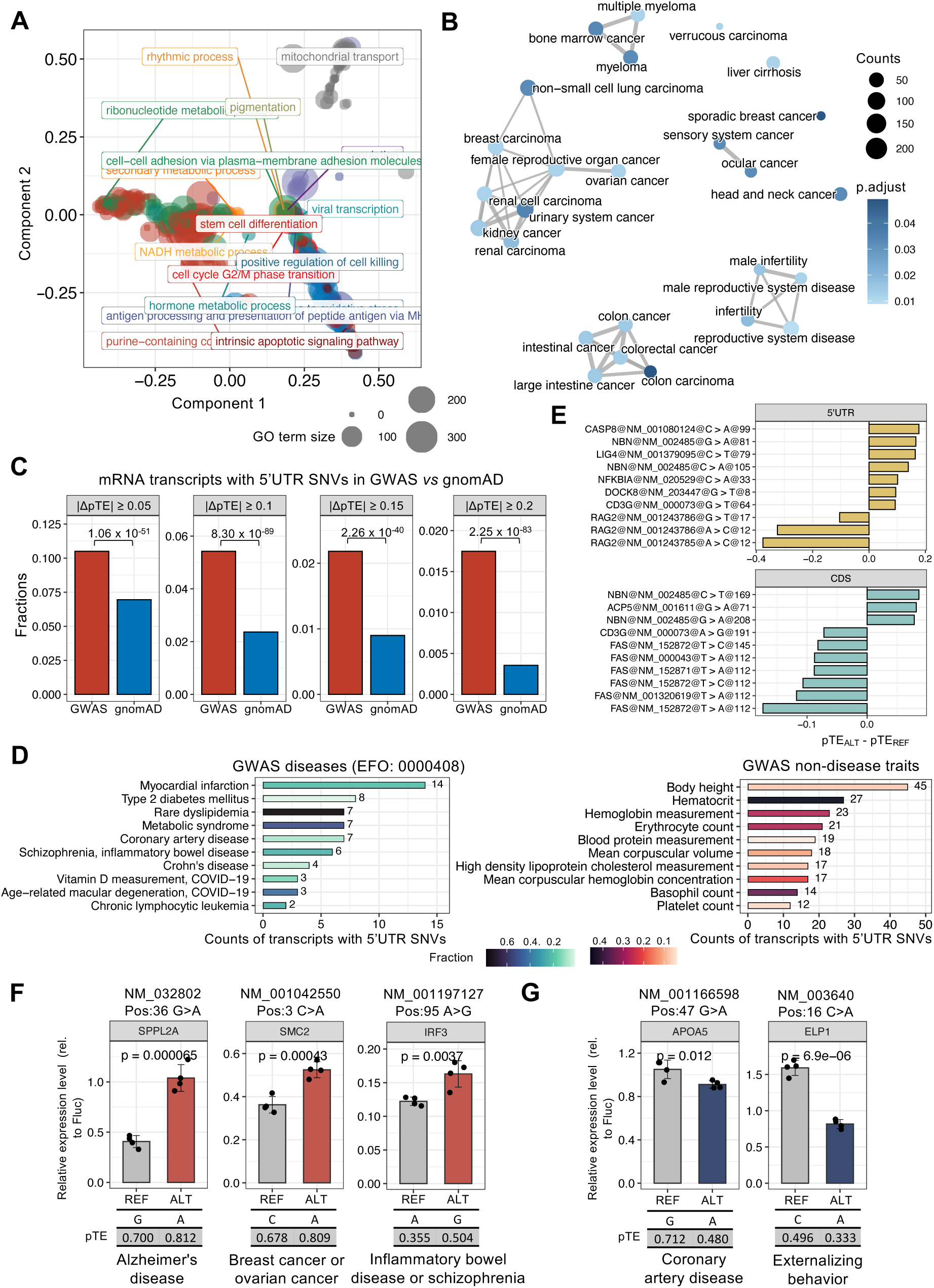
**Disease associations of genetic variants that alter translation.** (**A**) Construction and evaluation of processes for variants with |ΔpTE| > 0.1. Dot size indicates the number of genes the GO term contains; colors denote parent terms summarized by *rrvgo*. Axes represent the first two components from PCoA of the semantic similarity matrix. (**B**) Disease ontology enrichment for the same variant set as in (A). Dot size indicates gene counts; color scale shows adjusted p-values. (**C**) Fractions of transcripts with 5’UTR SNVs that passed various |ΔpTE| cutoffs in GWAS (red) and gnomAD (blue). P-values from Fisher’s exact test. (**D**) GWAS diseases (left) and non-disease traits (right) enriched for 5’UTR TE-altering SNVs (|ΔpTE| > 0.1). Bars show counts; color scale indicates the fraction of transcripts in each trait carrying such variants. The enriched traits with top 10 SNVs counts were filtered using p < 0.05 from Fisher’s Exact test. (**E**) Top 10 ClinVar immune-related SNVs ranked by |ΔpTE| in the 5’UTR (top) and CDS (bottom). (**F-G**) Luciferase reporter assays for GWAS disease-related SNVs. Bars show mean ± SD from biological triplicates; points indicate replicates; y-axis shows the normalized luciferase ratio. P-values from one-tailed t-tests are shown. TEFL-mRNA predicted pTE values for reference (REF) and alternative (ALT) alleles, along with associated GWAS traits, are listed below each plot.

We next examined whether disease- and trait-associated genetic variants from GWAS impact translation (Supplementary Table 4), and observed that a significantly larger fraction of GWAS 5’UTR SNVs altered TE compared to background variants in gnomAD. This enrichment became more pronounced with increasing |ΔpTE| thresholds, indicating that stronger TE effects are more prevalent among GWAS hits (**Figure 6C**). Further trait-specific analysis revealed associations between TE-altering SNVs and a diverse range of phenotypes, including cardiometabolic, immune, and hematologic-related traits (**Figure 6D**). Strikingly, all 13 GWAS missense variants that altered TE were predicted to decrease translation, in contrast to the more balanced distribution of TE changes observed among gnomAD missense variants (Supplementary Tables 1 **and 4**).

Focusing on ClinVar-annotated immune-related variants (particularly relevant given the use of LCL data in our model), we found that the top 10 SNVs with the largest |ΔpTE| (in 5’UTR or CDS regions) resided in key genes in immune signaling (Figure 6E). These include *RAG2*, involved in the V(D)J recombination during B and T lymphocyte development^69^; *NFKBIA*, a regulator of the NF-κB pathway essential for immune homeostasis^70^; *FAS* and *FASLG*, both central to apoptotic signaling and implicated in autoimmune lymphoproliferative syndrome^71^; and *CD3G,* a T cell receptor subunit linked to immune dysregulation^72^. These findings suggest that translational control by SNVs may play a critical role in shaping immune function and disease susceptibility (Supplementary Table 5).

To functionally validate disease-associated TE-altering SNVs, we also performed luciferase reporter assays (Figure 3G) on five GWAS variants. Three 5’UTR variants in *SUPPL2A*, *SMC2*, and *IRF3* were experimentally confirmed to enhance translation, reported as GWAS variants for Alzheimer’s disease, breast cancer, and inflammatory bowel disease or schizophrenia, respectively (Figure 6F). Two additional variants in *APOA5* and *ELP1* that we found to repress translation (Figure 6G) were linked to coronary artery disease and externalizing behavior in GWAS. Notably, these genes span multiple functional categories, including immune regulation (*SPPL2A*, *IRF3*), chromatin and cell cycle control (*SMC2*), lipid metabolism (*APOA5*), and neuronal function (*ELP1*), reflecting the broad impact of TE-altering SNVs on diverse biological pathways.

### Cross-cell-line generalization and disease relevance of translation-altering variants

To investigate whether TE-altering SNVs are cell-type-specific, we extended TEFL-mRNA to nine additional data sets from various cell types (one LCL replicate and eight other cell types). A separate model was trained per dataset and performed robustly (**Figure 7A**), indicating good generalizability across transcriptomes. We then predicted the impact of ExAC SNVs on TE, using a relaxed |ΔpTE| threshold (>0.02) to capture a wider spectrum of effects (corresponding to an estimated false positive rate < 1% in all datasets, Supplementary Figure 8A, Supplementary Table 6). Consistent with the LCL analysis (**Figure 3D**), TE-altering variants were most abundant in the 5’UTR and CDS (Supplementary Figure 8B).

**Figure 7.**
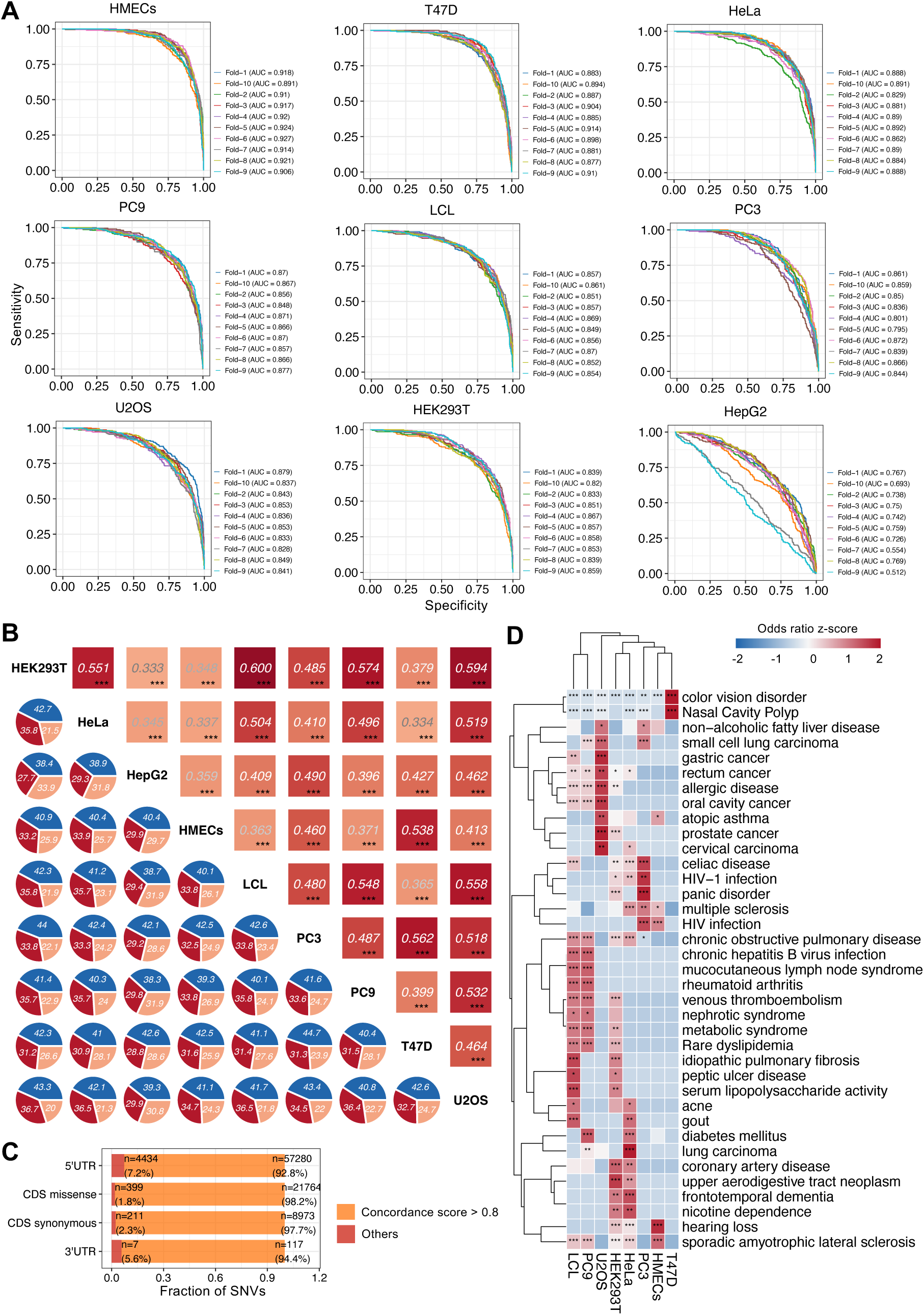
**Translation-altering SNVs across multiple cell types.** (**A**) Receiver-operating characteristic (ROC) curves for TEFL-mRNA models trained separately in nine human cell types. Performance was assessed by 10-fold cross-validation for the TE-high vs others classification; each fold is shown, and the per-model AUC is annotated. (**B**) Cross-cell-type agreement for 5’UTR SNVs. Tiles show the Spearman correlations of ΔpTE between each pair of cell types, computed on ExAC 5’UTR SNVs with |ΔpTE|>0.02 in both cell types (two-sided test; *** p < 0.001). Overlaid pie charts indicate the proportion of shared SNVs with concordant positive (red), concordant negative (blue), or discordant (salmon) signs of TE effects. (**C**) Sharing of TE effects across cell lines by region. Stacked bars give the percentage of SNVs that are “shared” across cell types (orange), defined by a concordance score >0.8 (the fraction of profiled cell types in which the ΔpTE sign matches the majority sign among cell types where |ΔpTE|>0.02), for 5’UTR, CDS missense, CDS synonymous, and 3’UTR SNVs. Count and percentage of SNVs contributing to each bar are shown. (**D**) Disease enrichment of TE-altering GWAS 5’UTR SNVs across cell types. Heatmap shows the enrichment of TE-altering SNVs (|ΔpTE|>0.02) associated with disease traits (EFO:0000408) in at least two cell types using Fisher’s exact test (p < 0.05); values are z-scores of the odds ratio in each trait across cell types. Row/column dendrograms indicate hierarchical clustering of traits and cell types (*** p<0.001, ** p<0.01, * p<0.05).

We further focused on the 5’UTR SNVs. Across pairs of cell lines, ΔpTE effects were significantly correlated in all cell line pairs, with 70-80% of variants displaying the same direction of effect (**Figure 7B**). To summarize concordance across all lines, we calculated a “concordance score” for each variant as the fraction of cell lines in which its ΔpTE direction matched the majority TE-altering directions among all cell lines. Over 90% of SNVs had scores >0.8, indicating highly reproducible directionality of TE effects across cell types (**Figure 7C**).

Consistent with the LCL results (**Figure 3**), pooling all cell lines showed that the TE-increasing alleles of 5’UTR SNVs were enriched for A/U alleles, located near the start codon, and skewed toward ultra-rare allele frequencies (Supplementary Figure 8C-E). Among GWAS SNVs, TE-altering 5’UTR SNVs were significantly overrepresented at loci associated with multiple types of cancer, immune, cardiometabolic, and neurodegenerative disorders (**Figure 7D**, Supplementary Table 7). While many enrichments were shared across cell types, the associations varied by cellular context, with cancer-related enrichments generally stronger in cancer-derived lines. These findings highlight both shared and cell-type–specific disease relevance of translation-altering 5’UTR variants.

We next tested whether CDS signals observed in LCLs generalize across cell types. Synonymous SNVs that increased TE did not necessarily introduce CAI-“optimal” codons (Supplementary Figure 8F), recapitulating the LCL result (**Figure 4C**). Missense variants encoding proline residues frequently reduced TE in 5 of the 9 cell lines (Supplementary Figure 8G). Notably, the direction of effect for GWAS missense variants was cell-type dependent: they skewed toward TE-decreasing in LCLs, U2OS, and PC9 but toward TE-increasing in the other cell lines (Supplementary Figure 8H).

Together, these results suggest a two-layer architecture of translational regulation by genetic variation: a conserved, broadly shared 5’UTR component, and a cell-type-dependent coding component where synonymous and missense effects vary in magnitude and direction, plausibly reflecting differences in codon-usage programs and tRNA supply across cell types.

## Discussion

Our study reveals translation as a widespread and underappreciated target of genetic variation, with tens of thousands of SNVs imposing measurable effects on protein output. While the impact of genetic variants on transcription and splicing has been extensively characterized, their influence on translation efficiency remains poorly defined. By systematically quantifying SNV effects at single-nucleotide resolution, we demonstrate that RNA translation is a major regulatory layer through which genetic variation shapes phenotypic outcomes.

A key finding is that missense variants, traditionally known only for their protein-recoding capacity, also exert significant and amino acid-specific effects on translation (Supplementary Figure 6C). In LCLs, substitutions introducing phenylalanine consistently increase TE, in agreement with the proposed role of tryptophan (W) to phenylalanine (F) substitutions in altering protein synthesis to activate T cell responses in tumor immunoreactivity^73^. In contrast, the substitutions introducing proline consistently reduced TE, with longer ones causing progressively larger decreases, supporting the ribosome stalling mechanism^74^. Extending this analysis to additional cell types, we found that proline-encoding missense variants reduced TE in five of the nine cell lines, indicating that this effect is reproducible but not universal. More broadly, the direction of TE changes for missense GWAS variants was strongly cell type-dependent: they skewed toward TE-decreasing in LCLs, U2OS, and PC9, but toward TE-increasing in other cell types. Synonymous variants also revealed unexpected behavior, with TE-increasing alleles not necessarily corresponding to codon “optimality,” a paradoxical effect reproduced across multiple cell types.

These coding variant observations are consistent with global sequence and structural correlates of TE. In LCLs, TE was negatively correlated with CAI and GC content, and positively correlated with AMFE. This pattern was shared by several other lines (HMECs, T47D, PC3, PC9, HeLa), but reversed in HEK293T, U2OS, and HepG2, where CAI and GC content correlated positively with TE (Supplementary Figure 9A-B). These results indicate that the relationship between codon usage, RNA structure, and translation efficiency is highly cell-specific. In some contexts, excessive codon optimization or stable GC-rich structures may impede translation^75,76^, while in others they may enhance efficiency, likely reflecting differences in ribosome abundance, tRNA pools, or codon usage programs. Such diversity highlights the complexity of translational regulation. Future work incorporating ribosome dynamics and cellular context will be essential for understanding and ultimately controlling translation in a cell type-specific manner.

In contrast to the cell type-specificity of coding variants, 5’UTR variants, especially A/U-biased SNVs near start codons, showed remarkably consistent TE effects across nine cell types, reinforcing their conserved role in regulating initiation. This two-layer regulatory architecture, conserved initiation control versus context-specific coding influences, has important implications for variant interpretation and for understanding tissue-specific mechanisms of genetic risk.

The disease relevance of TE-altering variants further highlights their importance. Variants altering TE were enriched in immune and cancer pathways and overrepresented among GWAS loci, with 5’UTR SNVs showing the most consistent disease associations across cell types. Coding variants, particularly GWAS loci linked with missense substitutions, demonstrated cell type-dependent effects, highlighting translational control as a mechanistic link between genetic variation and complex traits.

Finally, our framework provides a scalable discovery engine for functional variant interpretation. TEFL-mRNA not only recapitulated known regulatory features such as uORFs and miRNAs but also uncovered a granular view of codon usage, RNA secondary structures, and RBP binding. While current models remain limited by incomplete structural and ribosome dynamics data, the strong reproducibility of 5’UTR variant effects across cell types and the novel insights into coding variants highlight the robustness of sequence-based inference. Looking forward, integration with ribosome profiling in primary tissues, tRNA landscapes, and generative modeling approaches could enable precise design of mRNAs with tunable translational properties.

In conclusion, our work establishes translation as a pervasive mechanism through which genetic variation influences protein output and disease risk. By systematically mapping translation-altering variants across the genome and across cell types, we uncover a previously hidden layer of gene regulation and provide a framework for functional variant prioritization in complex traits and diseases.

## Methods

### Ribosome profiling data processing and translation efficiency evaluation

Ribosome profiling and matched RNA-seq data in lymphoblastoid cell lines derived from 72 HapMap Yoruba individuals in Ibadan, Nigeria (YRI) were downloaded from GEO (accession GSE61742 and GSE19480)^25,77^. Quality control and adaptor trimming were performed using FastQC (version 0.11.9, https://www.bioinformatics.babraham.ac.uk/projects/fastqc/) and Trim Galore (version 0.6.6, https://github.com/FelixKrueger/TrimGalore). Bowtie2 (version 2.4.2) was used to remove reads mapped to ribosomal RNA^77^. Then, clean reads were mapped to the human reference genome (GRCh38) using STAR (version 2.7.6)^78^. Gene expression quantification was performed with RSEM^79^. Translation efficiency (TE) of each transcript was calculated by dividing the TPM from Ribo-seq by the TPM from RNA-seq.

From the initial set of 63,748 curated mRNAs from RefSeq, we first filtered out mRNAs based on high missingness (TE value missing in > 50% samples) and low expression (average RNA-seq read counts ≤10), which retained 15,844 mRNAs for further analysis. To ensure consistency across samples, we calculated pair-wise correlations of transcript-level TE values and retained 47 samples with average correlation coefficients above 0.7. After further excluding the mRNAs with missing TE values in more than 5 samples, 9,745 mRNAs remained for downstream classification. Transcripts were categorized as highly translated (“TE-high”), lowly translated (“TE-low”), and intermediate (“TE-intermediate”) in each sample using the cutoffs TE > 1, TE < 0.25, and 0.25 <= TE <= 1, respectively. To select consistently classified transcripts, we labeled mRNAs as TE-high or TE-low if they met the respective criteria in more than 75% of samples. To ensure balanced class sizes for neural network training, we retained only those transcripts classified as TE-intermediate in more than 90% of samples. This resulted in 726 TE-high, 524 TE-low, and 729 TE-intermediate mRNAs. Full-length mRNA sequences and coding region annotations were extracted from the NCBI RefSeq database.

### Sequence embedding of full-length RNA sequences

To accommodate variable mRNA lengths with the fixed input dimensions required by the CNN, we padded all sequences to match the maximum transcript length observed (19,940 nucleotides). Then, each mRNA sequence was represented by one-hot encoding, which generates a binary matrix of size 4 x 19,940, where rows correspond to A, G, C, and U. In addition, we added a fifth row indicating the mRNA regions (‘0’ for 5’UTR; ‘1’ for CDS; ‘2’ for 3’UTR). Finally, all mRNA sequences were encoded into a unified input matrix with dimensions (N, 19940, 5), where N was the total number of transcripts.

### Convolutional and recurrent hybrid neural network

Deep learning models, particularly CNNs and RNNs, have demonstrated remarkable success in various sequence analysis tasks, including DNA motif discovery, protein structure prediction, and RNA secondary structure prediction^30^. CNNs are adept at recognizing spatial patterns in data, making them well-suited for identifying sequence motifs, while RNNs excel at modeling sequential data, capturing long-range dependencies crucial for understanding RNA regulatory mechanisms. TEFL-mRNA integrates CNNs and Bidirectional Long Short-Term Memory (BiLSTM) layers to effectively identify both proximal and distal features within full-length mRNA sequences.

The CNN component extracted spatial features and sequence motifs potentially involved in translational regulation^80^. The architecture included several convolutional layers. First, the input layer took the one-hot encoded RNA sequence matrix, followed by multiple convolutional layers with varying kernel sizes to capture diverse local motifs within the sequences. Each convolutional layer was followed by a Rectified Linear Unit (ReLU) activation function. Next, max-pooling layers were employed to reduce the dimensionality of the feature maps, retaining the most significant features and reducing computational complexity. Dropout layers were incorporated to mitigate overfitting. This structure allowed the CNN to learn and identify important sequence motifs that may function as *cis*-regulatory elements.

The RNN component employed BiLSTM layers to model sequential dependencies and long-range interactions. LSTMs are particularly effective in managing long-range dependencies and mitigating issues such as vanishing gradients in traditional RNNs^81^. The LSTM cell usually consists of three gates: forget gate, input gate, and output gate,

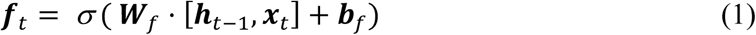

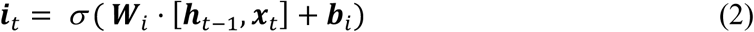

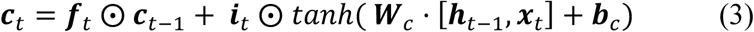

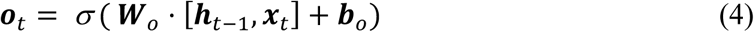

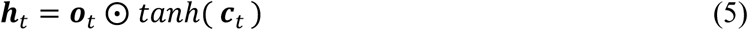

where σ denotes the sigmoid activation function, with output values between 0 and 1. *W_f_*, *W_i_*, *W_c_*, and *W_o_*_’_ are the weight matrices for the forget, input, cell, and output gates, respectively. *b_f_*, *b_i_*, *b_c_*, and *b_o_*_’_ represent the bias terms for respective gates. *h_t-_*_1_ is the hidden state from the previous time step. *x_t_* is the input at the current time step and :*c_t_* is the cell state, which carries information across different time steps.

The BiLSTM layers were followed by dropout layers to further prevent overfitting. Output from these layers was flattened and passed through fully connected (dense) layers to perform the final feature combination and transformation, enabling the model to make predictions about RNA sequence functions. Finally, an output layer with a SoftMax activation function was used for classification, providing probability distributions over the possible RNA functions. This integrated model architecture allowed us to combine the strengths of both CNNs and RNNs, enabling the model to capture both local and global regulatory features within the RNA sequences.

### Model performance was evaluated using accuracy, precision, recall, F1-score, and ROC-AUC metrics

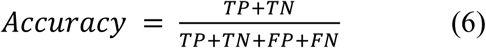

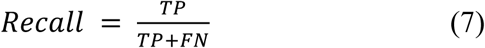

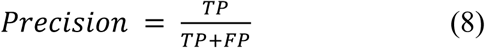

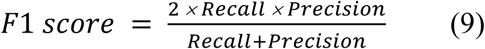

where TP, TN, FP, and FN indicate the number of true positives, true negatives, false positives, and false negatives, respectively. Moreover, we used the ROC (receiver operating characteristic) curve to intuitively evaluate classification performance.

Hyperparameter tuning was performed using the Tree Parzen Estimator (TPE) algorithm^82^, optimizing parameters including batch size, kernel size, filter size, optimizer, and dropout rates across 100 iterations. The optimized parameters include the batch size of 128, filter size of 64, kernel size of 11, as well as three dropout rates of 0.8873, 0.2516, and 0.6746, respectively. The model was trained for a maximum of 2,000 epochs, using a categorical cross-entropy loss function and the RMSprop optimizer. In addition, we employed multiple strategies to reduce over-fitting, including batch normalization^83^, dropout^84^, and early stopping based on validation loss. In total, 928,771 parameters were trained in our model (Supplementary Figure 1). TEFL-mRNA was implemented using TensorFlow and Keras libraries. The training and evaluation were performed on a high-performance computing cluster with GPU acceleration (NVIDIA A100 and H100 GPUs).

### Upstream open reading frame and microRNA binding prediction

To evaluate whether our model captured established mechanisms of translational regulation, particularly translation inhibition mediated by uORF and miRNA binding sites, we compared predicted pTE between mRNAs with and without these regulatory elements. Human uORF annotations were downloaded from uORFdb, which includes >2.4 million human uORFs^85^. We selected mRNAs annotated with “strong” uORFs as the ‘with uORFs’ group. For miRNA binding predictions, we utilized miRWalk^86^, selecting mRNAs with miRNA binding probability higher than 0.9 in the 3’UTR or 5’UTR. To further refine miRNA targets relevant to our cellular context, we integrated miRNA expression data from naïve B cells obtained from miRbase, selecting mRNAs targeted by at least one expressed miRNA as the “with miRNA binding sites” group.

### Calculation of sequence and structural features and binning analysis

We calculated a series of sequence and structure features for each full-length mRNA. RNA secondary structures were predicted using LinearFold with the --verbose parameter^87^. The estimated half-life and RNA degradation scores were evaluated using the DegScore model, a ridge regression-based method trained on Eterna Roll-Your-Own structure data for RNA degradation prediction (https://github.com/eternagame/DegScore)^88^. The Minimum Free Energy (MFE) values and harpin counts were extracted directly from the LinearFold results using the custom scripts. Adjusted MFE (AMFE) was computed by dividing the MFE by mRNA length. We also calculated CAI, a measure of the codon usage adaptiveness introduced by Sharp and Li in 1987^89^, defined as follows:

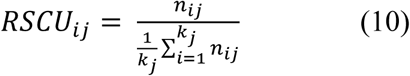

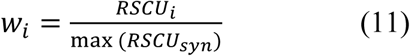

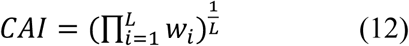

where *n_ij_* is the number of occurrences of the *i_th_* codon encoding the *j_th_* amino acid.*k_j_* represents the number of synonymous codons for the *j_th_* amino acid. The relative adaptiveness *w_i_* for each codon was calculated by normalizing its Relative Synonymous Codon Usage (RSCU) value by the maximum RSCU among synonymous codons of the same amino acid. Finally, for an mRNA containing L codons, CAI was calculated as the geometric mean of the *w_i_* values for all the codons in the gene.

### Human genetic variants data collection, preprocessing, and annotation

We compiled SNVs from two large-scale datasets: the ExAC database (60,706 exomes) and gnomAD v4.0 (730,947 exomes; https://gnomad.broadinstitute.org). Variants located in protein-coding genes and curated RefSeq mRNAs were extracted. For downstream analyses, we focused on SNVs annotated as ‘5_prime_UTR_variant’, ‘3_prime_UTR_variant’, ‘synonymous_variant’, and ‘missense_variant’ according to the Variant Effect Predictor (VEP)^90^. In total, 2,634,150, 4,261,376, 5,397,223, and 6,044,404 unique variants were categorized as 5’UTR, 3’UTR, synonymous, and missense variants, respectively. Common variants (as controls) were extracted from dbSNP (https://www.ncbi.nlm.nih.gov/snp/). Additionally, we obtained uAUG-creating variants previously reported to affect translation initiation^56^, ClinVar variants (https://www.ncbi.nlm.nih.gov/clinvar/), and trait-associated variants from the GWAS catalog (v1.0.2; https://www.ebi.ac.uk/gwas/home). The TE effect of each genetic variant was defined as pTE_alt_ – pTE_ref_ (ΔpTE)

### Prediction of mRNA structures and RBP binding changes induced by SNVs

RNA secondary structures harboring the reference and alternative alleles were predicted using LinearFold and full-length mRNA sequences^87^. To assess changes in RBP affinity, we used DeepBind^66^, a deep learning-based tool for RBP binding prediction. For each SNV, we extracted a 51-nucleotide (nt) window centered on the variant (25nt upstream and downstream), generating two sequences – one with the reference allele and one with the alternative allele. Binding scores for each allele were calculated using DeepBind, and the difference in scores (ΔRBP_BS) was calculated as ΔRBP_BS = RBP_BS_Alt_ – RBP_BS_Ref_.

To identify RBPs with potential regulatory roles, we compared ΔRBP_BS values between TE-altering variants (|ΔpTE| > 0.1) and TE-unchanged variants (defined as |ΔpTE| < 10^-6^), using the Student’s t-test. RBPs with significant differences (p < 0.05) were considered putative regulators of translation.

To assess positional effects of SNVs within RBP motifs, we focused on variants associated with the top and bottom 5% of ΔRBP_BS values as TE-increasing and TE-decreasing SNVs, respectively. We applied a hexamer sliding window across the 51nt sequences. Each hexamer was scored against RBP position weight matrices (PWMs) using a normalized scoring function (scaled to [0, 1]) based on nucleotide frequency weights. We identified specific motif positions where SNVs most strongly disrupted or enhanced RBP binding (highest score changes). The PWMs were downloaded from cisBP-RNA^91^ (catalogue of inferred sequence binding preferences for RNA) (http://cisbp-rna.ccbr.utoronto.ca/).

### Functional enrichment and disease relevance analysis of the TE-altered genetic variants

We defined TE-altering genetic variants as those with absolute changes in pTE higher than 0.1 (|pTE_alt_ - pTE_ref_| > 0.1). Gene ontology (GO) and disease ontology (DO) enrichment analyses were conducted using *clusterProfiler*^92^ and *DOSE*^93^, with the adjusted p-value threshold of <0.05 for significance. To group the GO terms for further visualization, we used the R package *‘rrvgo’* to calculate the semantic similarity based on biological process annotations. GO terms were grouped using a similarity threshold of 0.9^94^, and representative parent terms were extracted. Visualization of enriched GO terms was generated using the *scatterplot* function, and the enriched DO terms were visualized with the *emapplot* function. GWAS traits were categorized into disease and non-disease traits, with disease traits defined as descendants of EFO:0000408 using the R package ‘*gwasrapidd*’^95^. The enrichment of SNVs in each GWAS trait was evaluated using Fisher’s Exact test. Reference alleles were retrieved from the genome using Bedtools getfasta^96^. For functional analyses, we retained SNVs located in 5’UTR and CDS that mapped to curated RefSeq mRNAs.

### Cell type-specific TE prediction and variant-effect analysis

TE measurements for multiple human cell lines were obtained from Zheng et al^97^ . We selected 9 widely used cell lines: HEK293T, HeLa, HepG2, HMECs, LCL, PC3, PC9, T47D, and U2OS. Within each dataset, transcripts were classified into TE-high, TE-intermediate, and TE-low groups using the 75th and 25th percentiles of measured TE as thresholds. For each cell line, we trained a separate TEFL-mRNA model (same architecture as described above) on full-length mRNA isoforms encoded with region labels. Hyperparameters were tuned once using the average TE across cell types and then fixed for all models (batch size = 256, filter size = 128, kernel size = 7, dropout rates = 0.7418/0.4394/0.2471). Models were optimized with categorical cross-entropy loss and the Adam optimizer and evaluated by 10-fold cross-validation. The trained model produced, for each isoform, a probability of being TE-high (pTE). To quantify variant effects, we generated paired sequences carrying the reference and alternative allele for each SNV from three sources: dbSNP common variants (allele frequency 0.1-0.9), ExAC, and the GWAS catalog. For every variant-isoform pair, we scored pTE for the reference (pTE_ref_) and alternative (pTE_alt_) sequences and defined the effect size as ΔpTE = pTE_alt_ - pTE_ref_.

### Plasmid construction and transfection

To evaluate the impact of 5’UTR variants on translation efficiency, we synthesized variant-containing 5’UTR sequences (GENEWIZ) and cloned them into the Fluc-Nluc dual luciferase reporter vector^98^ (**Figure 3G**). The construct consists of a firefly luciferase (Fluc) gene driven by a constitutive promoter, with the test sequence harboring the SNV inserted into the 5’UTR. A NanoLuc (Nluc) luciferase gene served as an internal control for normalization. GM12878 cell line was cultured in RPMI 1640 (Gibco) medium containing 15% fetal bovine serum (FBS, Hyclone). To transiently transfect plasmids into cells, 1 μg of reporters were transfected into 6*10^5^ cells by using the 4D-Nucleofector (Lonza) in SF media (Lonza) with the DN-100 program^99^. Cells were incubated for 30 hours at 37 ℃, then collected for further analysis.

### RNA extraction and RT-PCR

Total RNA was extracted using the EasyPure Fast Cell RNA Kit (TransGen) following the manufacturer’s protocol. One microgram of total RNA was reverse-transcribed into cDNA using the PrimeScript RT Reagent Kit (Takara). RNA levels were quantified by qPCR using 2X Universal SYBR Green Fast qPCR Mix (Abclonal). The following primers were used in the qPCR:

Nluc-fwd: 5’-GTTGGGGACTGGCGACAGAC-3’;

Nluc-rev: 5’- TCATCCACAGGGTACACCAC-3’;

Fluc-fwd: 5’- GTTCGTCACATCTCATCTACCTCC-3’;

Fluc-rev: 5’- TGTAGTAAACATTCCAAAACCGTG-3’

### Luciferase activity assay

Cells were lysed using the Passive Lysis Buffer (Promega), and luciferase activities were measured using the Nano-Glo Dual-Luciferase Reporter Assay System (Promega), following the manufacturer’s protocol. The final luciferase activity was calculated as the ratio of Nluc to Fluc activity, followed by normalization to RNA levels measured by qPCR. This normalized luciferase ratio was used to compare translation efficiency between reference (REF) and mutant (MUT) constructs. This design allowed for a direct assessment of the impact of SNVs on translation in a controlled experimental setting.

## Acknowledgments

The authors want to thank Dr. Jiefu Li, Dr. Yue Hu, and Hanwen Zhou in the Wang Lab, and Dr. Giovanni Quinones-Valdez, Ryo Yamamoto, Elaine Huang in the Xiao Lab for their discussions and comments.

## Funding

This work is supported by the Natural Science Foundation of China (32030064 and 32250013), the National Key Research and Development Program of China (2021YFA1300503), the BMGF-NSFC Joint Project on Vaccine Research and Development (W2412022 and 2024VTJP1001), and the CAS Strategic Priority Research Program (XDB38040100) to Z.W.

## Author contributions

S.W., Z.W., and X.X. conceived and designed the study. S.W. analyzed the data, built the deep learning model, and conducted the statistical analysis and experiments. C.C. designed and conducted the experimental validation. S.W., Z.W., and X.X wrote the manuscript. Z.W. and X.X. provided key suggestions for study design and statistical analysis.

## Competing interests

The authors declare that they have no competing interests.

## Data and materials availability

The ribosome profiling and RNA-seq data were obtained from Gene Expression Omnibus (accession numbers GSE61742 and GSE19480). Genetic variants were downloaded from the gnomAD (v4.0) and ExAC databases (https://gnomad.broadinstitute.org). Common variants were obtained from the dbSNP database (https://www.ncbi.nlm.nih.gov/snp/). Disease-related variants were downloaded from ClinVar (https://www.ncbi.nlm.nih.gov/clinvar/) and GWAS Catalog (https://www.ebi.ac.uk/gwas/home). TEFL-mRNA is available at GitHub (https://github.com/CikiyWang/TEFL-mRNA). All data are available in the main text or the supplementary materials.

## Notes

### Competing Interest Statement

The authors have declared no competing interest.

