## Supplementary figures for "Genetic variation shapes human mRNA translation and disease risk"

### Supplementary Figures and Figure legends

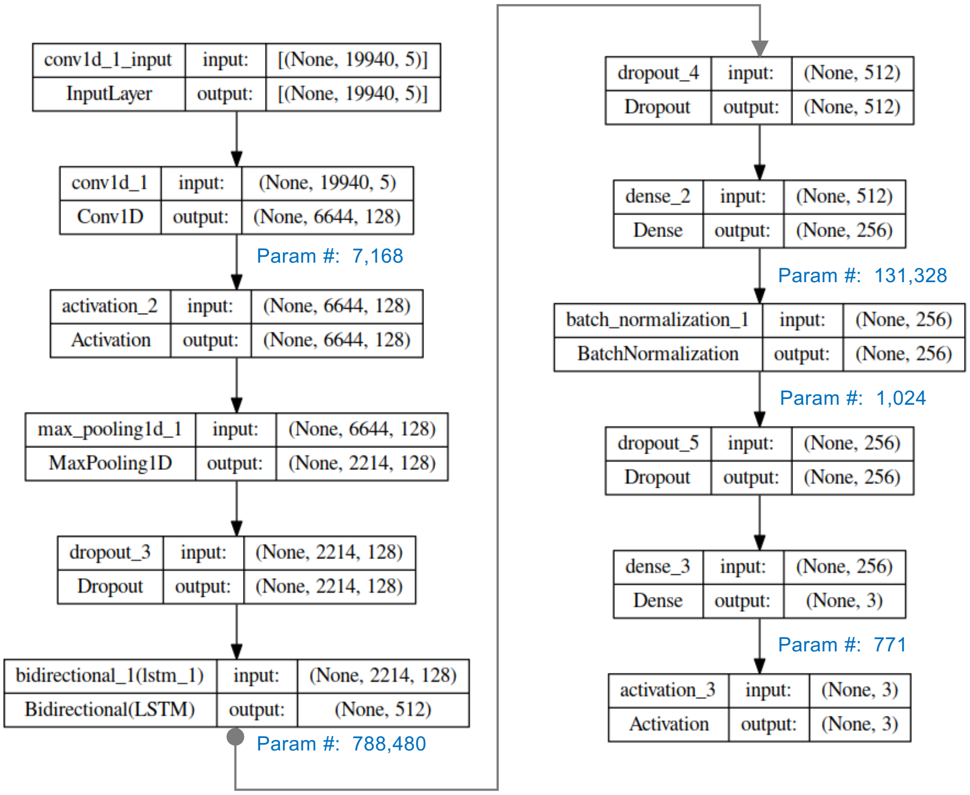


**Supplementary Figure 1. Architecture and performance of the TEFL-mRNA model for sequence-based TE classification.**

Schematic of the TEFL-mRNA architecture. Input tensor of shape (19940, 5) is processed through a 1D convolutional layer (Conv1D, 128 filters, kernel size 11), followed by activation and max pooling to reduce spatial dimensionality. A dropout layer provides regularization before a bidirectional LSTM layer captures long-range sequence dependencies in both forward and backward directions, producing a 512-dimensional feature vector. This is followed by additional dropout and two fully connected (Dense) layers. A batch normalization layer is applied between the dense layers to stabilize training. The final Softmax activation layer outputs probabilities for 3 TE classes. Parameter counts for each trainable layer are shown in blue. The model structure was visualized using *keras.utils.plot_model*.

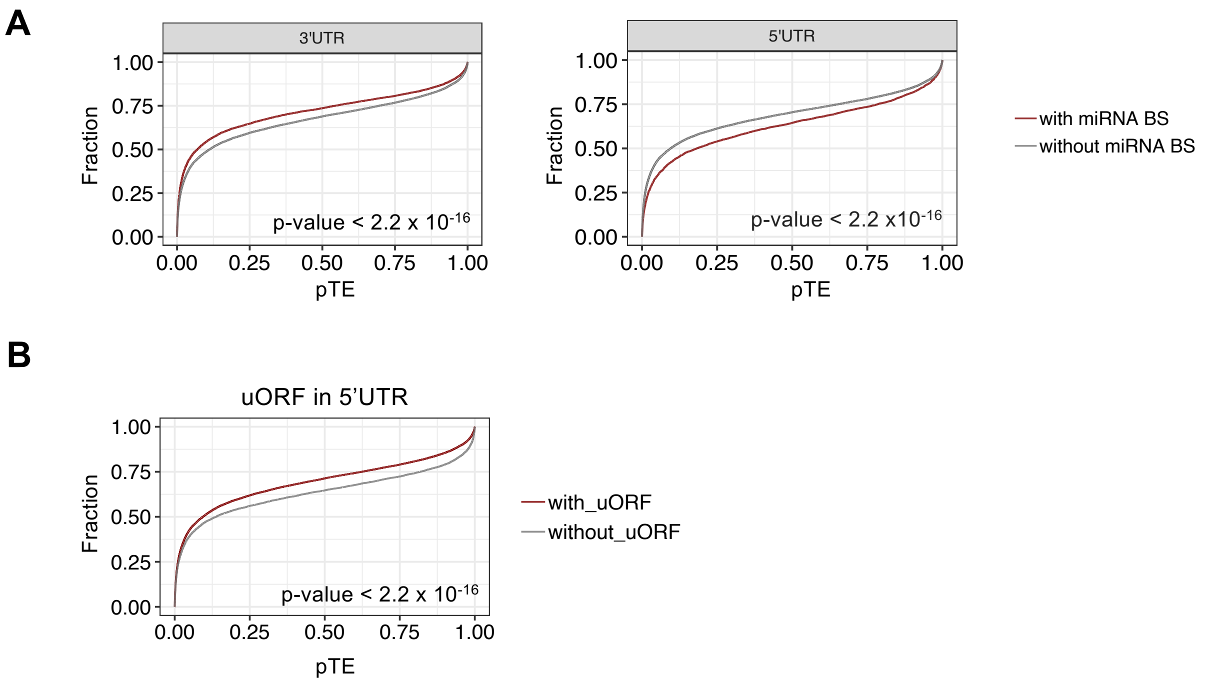


**Supplementary Figure 2. Prediction of the TE effects of miRNA binding and uORF.**

(**A**) Distribution of pTE for transcripts with versus without miRNA binding sites (BSs) (red versus grey lines) in the 3'UTR (left) or 5'UTR (right). P-values from Kolmogorov–Smirnov tests are shown.

(**B**) Distribution of pTE for transcripts with versus without uORF (red versus grey lines) in the 5'UTR. P-value from the Kolmogorov–Smirnov test is shown.

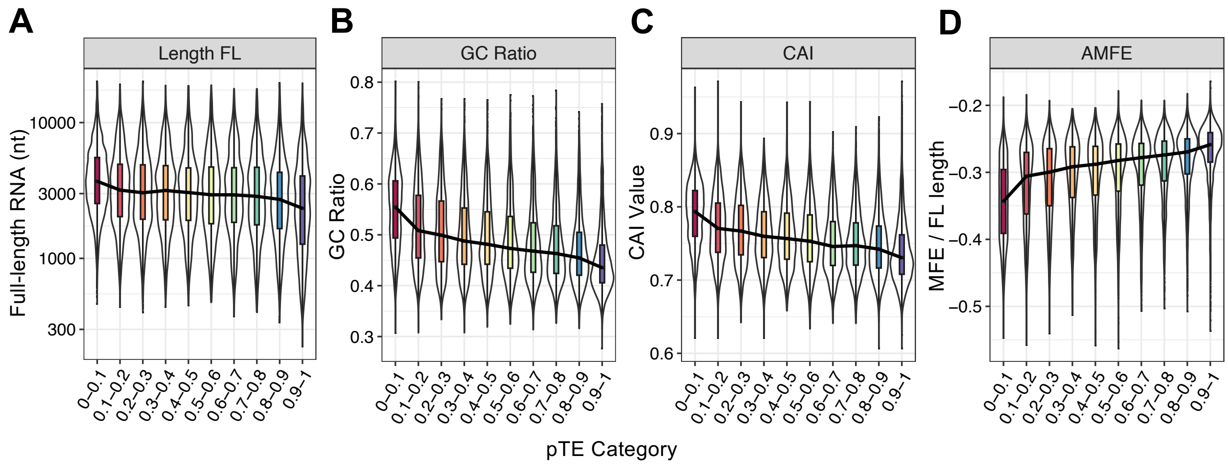


**Supplementary Figure 3. Relationship between sequence features and pTE categories.**

Violin plots showing the distribution of (**A**) mRNA length, (**B**) GC ratio, (**C**) CAI, and (**D**) adjusted MFE (AMFE) across pTE categories. Transcripts were divided into 10 bins based on the pTE values ranging from 0 to 1.

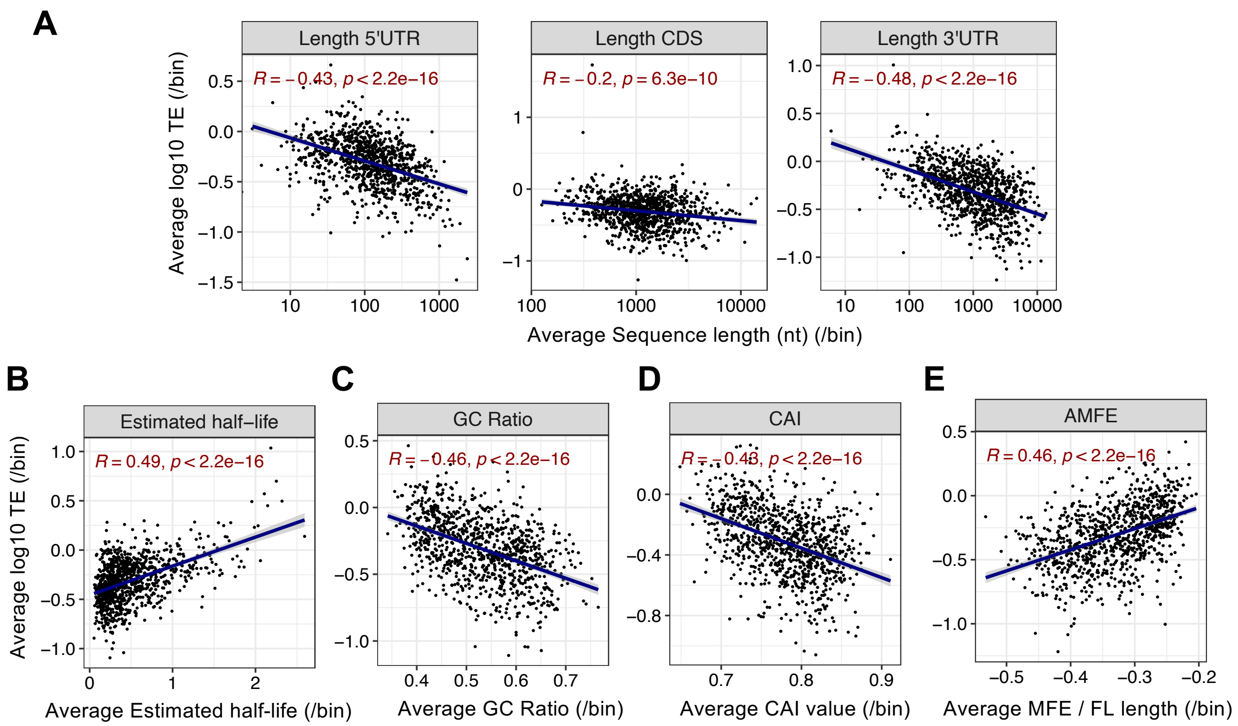


**Supplementary Figure 4. Sequence and structural features correlated with actual TE values.**

Scatter plots showing correlations between experimentally measured TE values versus known sequence or structural features. mRNAs were grouped into bins (10 mRNAs per bin), and axes show bin-averaged feature values versus bin-averaged TE. Pearson's Correlation coefficients and p-values are shown. (**A**) Lengths of the 5'UTR, CDS, and 3'UTR. (**B**) Estimated mRNA half-life. (**C**) GC ratio. (**D**) CAI. (**E**) AMFE.

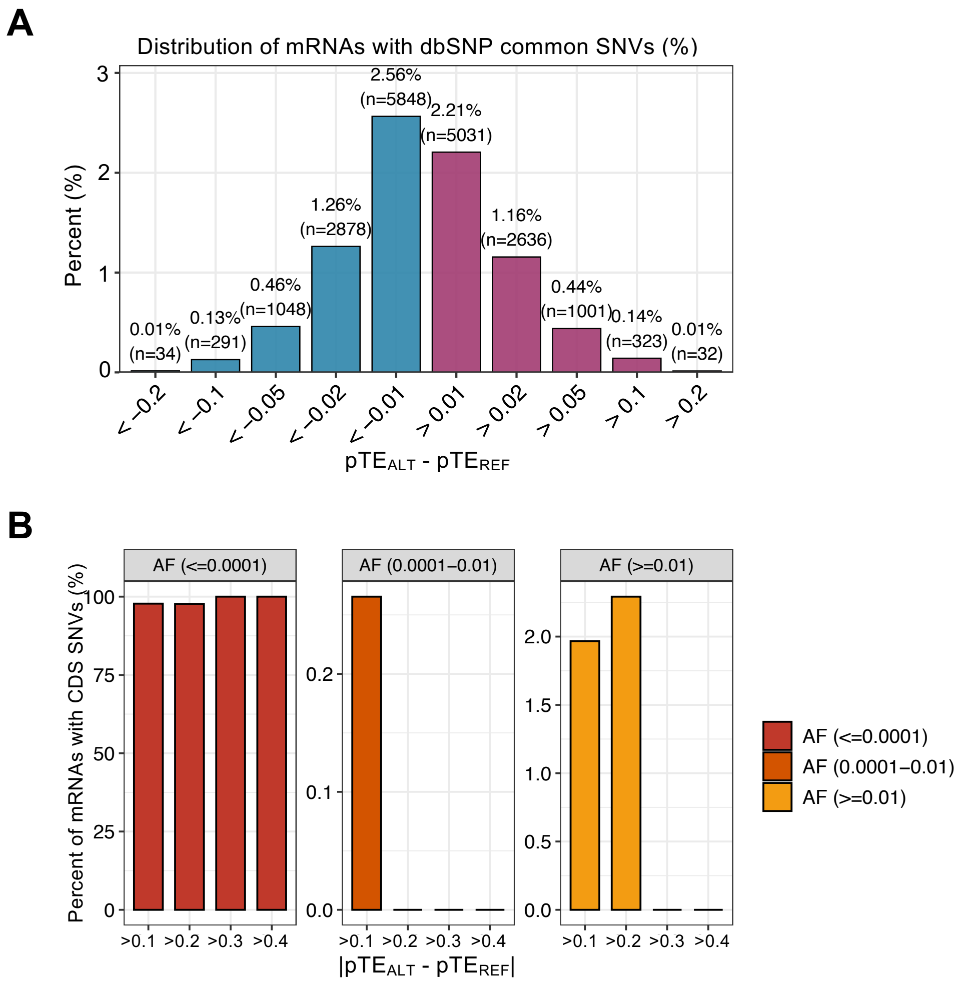


**Supplementary Figure 5. TE effects of dbSNP common variants and gnomAD CDS variants.**

**(A)** Percentage of dbSNP common (non-GWAS and non-ClinVar) variants (allele frequency between 0.1 and 0.9) passing different ΔpTE cutoffs. The y-axis shows the percent of SNVs passing specific ΔpTE cutoffs.

**(B)** Percentage of transcripts with gnomAD CDS variants passing the specified |ΔpTE| cutoff and within the allele frequency (AF) range among all transcripts with 5'UTR SNVs passing the |ΔpTE| cutoff.

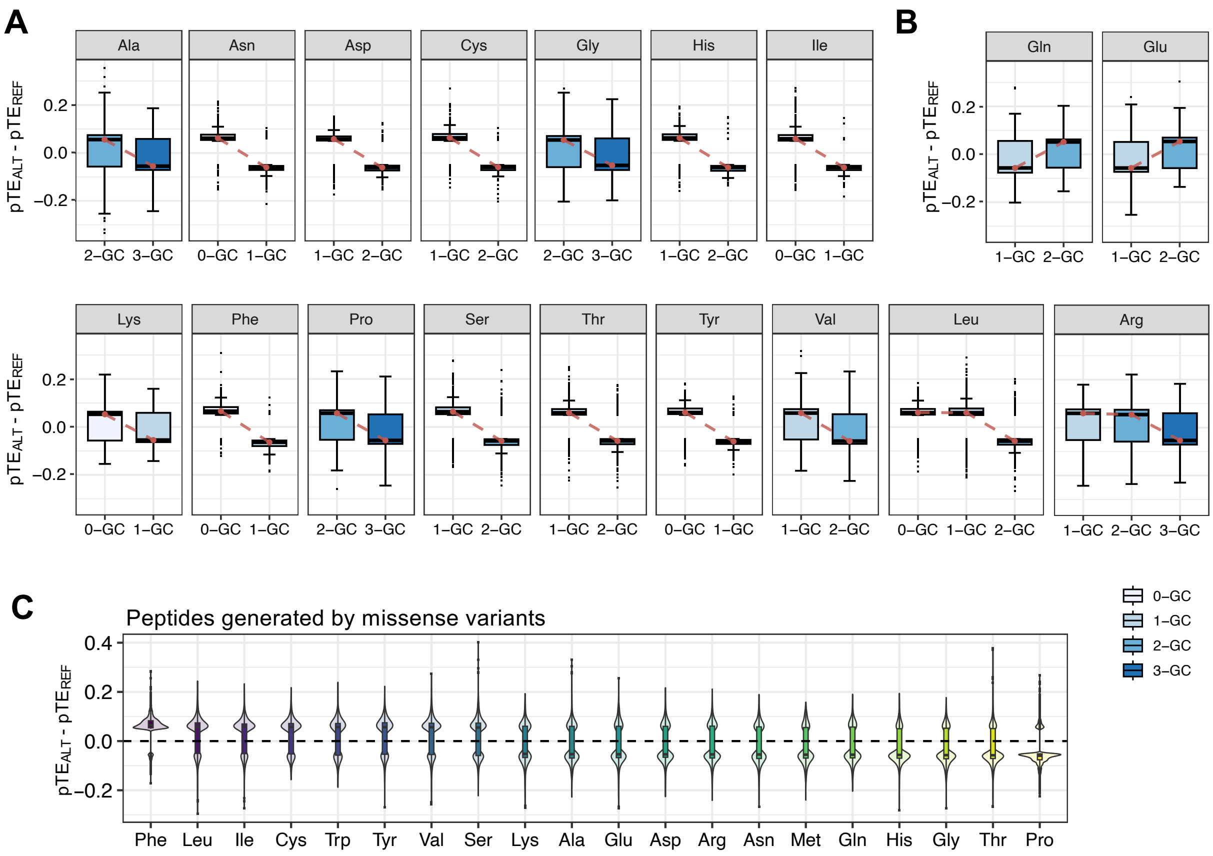


**Supplementary Figure 6. TE effects of SNVs in the CDS region.**

(**A-B**) Boxplots of ΔpTE values for synonymous SNVs with |ΔpTE| > 0.05, grouped by encoded amino acid. Within each amino acid, SNVs are classified by the number of G/C bases in the codons with alternative alleles (0-GC, 1-GC, 2-GC, and 3-GC).

(**C**) Violin plots showing ΔpTE distributions for missense SNVs that introduce different amino acids, restricted to variants with |ΔpTE| > 0.05. Codons are colored by the number of G/C bases in the codon with alternative alleles.

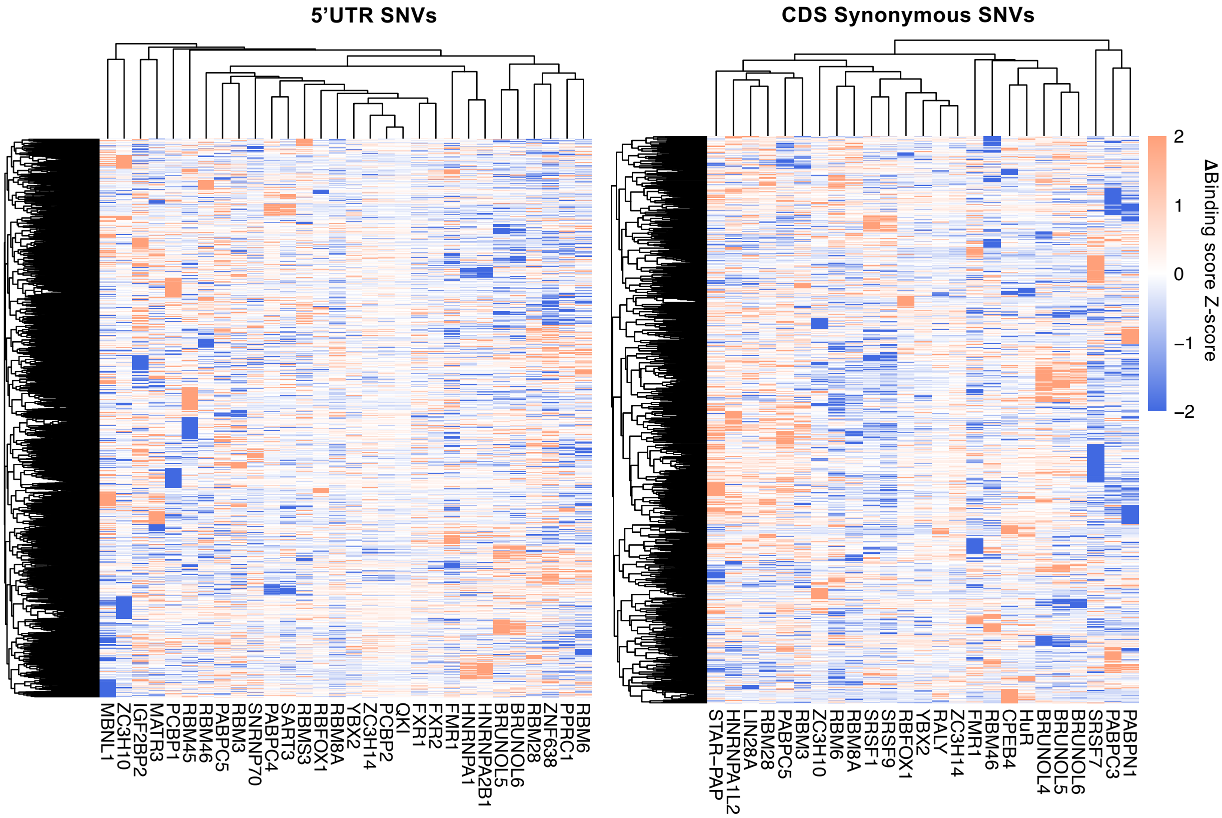


**Supplementary Figure 7. RBP binding perturbed by TE-altering SNVs.**

Heatmaps showing predicted changes of RBP binding scores (BS) for TE-altering SNVs in the 5'UTR (left) and CDS (synonymous SNVs; right). Binding score changes ΔBS were calculated using DeepBind on +/- 25nt sequences flanking each SNV, and Z-score-normalized by SNV (row). Blue indicates decreased binding; red indicates increased binding.

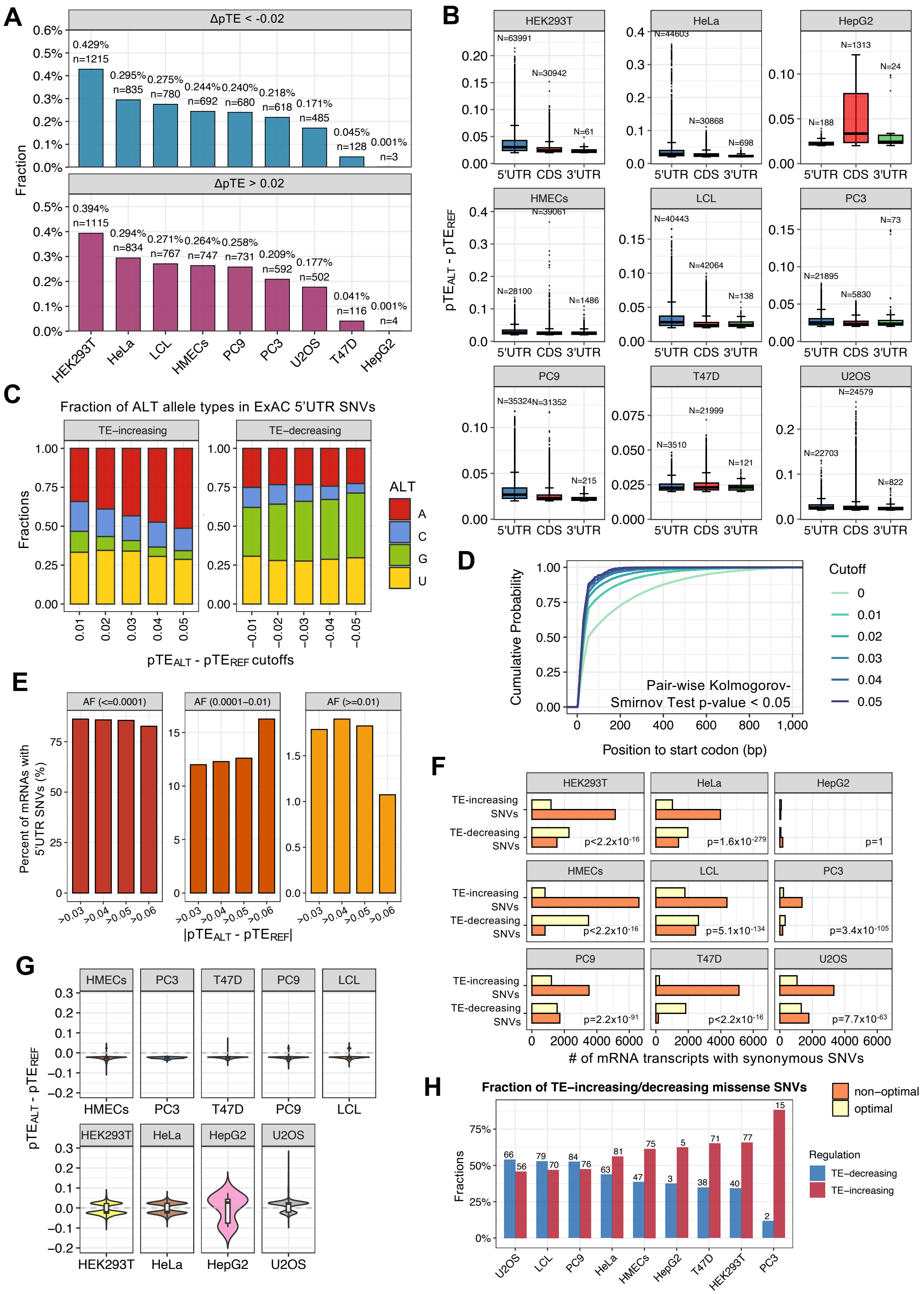

**Supplementary Figure 8. Translation effects of TE-altering variants across cell lines.**

**(A)** Fraction of common dbSNP SNVs (allele frequency 0.1-0.9) with ΔpTE below -0.02 (top) or above 0.02 (bottom panel) in each cell line.

(**B**) Regional distribution of TE-altering SNVs (|ΔpTE| > 0.02) in each cell line. Boxplots show ΔpTE; the number of SNVs contributing to each box is indicated.

**(C)** Alternative-allele composition of ExAC 5'UTR SNVs as stringency increases. Stacked bars give the proportions of alternative (ALT) allels (A/C/G/U) among TE-increasing (left) and TE-decreasing (right) SNVs passing the indicated ΔpTE cutoffs. The y-axis shows the fractions for 4 types of alternative alleles.

(**D**) Cumulative distribution of distances from 5'UTR SNVs to the start codon for increasing |ΔpTE| thresholds.

(**E**) Percentage of transcripts with ExAC 5'UTR SNVs passing the different |ΔpTE| cutoffs within the allele frequency (AF) range among all 5'UTR SNVs passing the |ΔpTE| cutoffs.

(**F**) Counts of TE-increasing and TE-decreasing synonymous SNVs across cell lines. Orange bars: variants changing codons toward non-optimal; yellow bars: toward optimal. P-value from Fisher's exact test.

(**G**) The distribution of ΔpTE of the TE-altering missense ExAC SNVs with |ΔpTE| > 0.02 that encode proline across 9 cell lines.

(**H**) The percentage of TE-increasing (red bars) and TE-decreasing (blue bars) missense GWAS SNVs across cell lines (|ΔpTE| > 0.02). The number of transcripts with missense SNVs in each cell line is labelled above each bar.

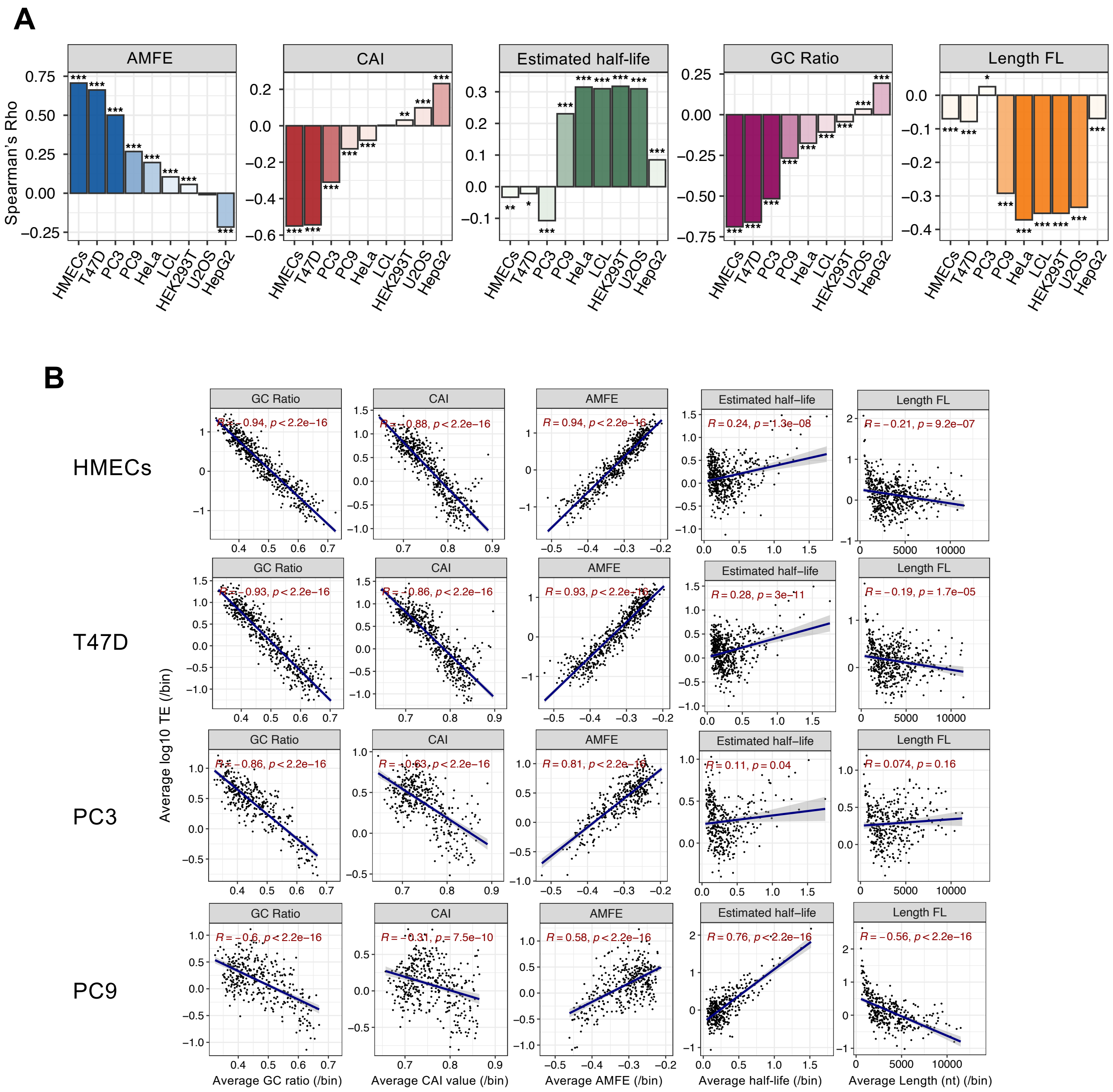
A
AMFE
CAI
Estimated half-life
GC Ratio
Length FL
Spearman's Rho
HMECs
T47D
PC3
PC9
HeLa
LCL
HEK293T
U2OS
HepG2
B
HMECs
T47D
PC3
PC9
Average log10 TE (/bin)
GC Ratio
CAI
AMFE
Estimated half−life
Length FL
R = −0.94, p < 2.2e−16
R = −0.88, p < 2.2e−16
R = 0.94, p < 2.2e−16
R = 0.24, p = 1.3e−08
R = −0.21, p = 9.2e−07
R = −0.93, p < 2.2e−16
R = −0.86, p < 2.2e−16
R = 0.93, p < 2.2e−16
R = 0.28, p = 3e−11
R = −0.19, p = 1.7e−05
R = −0.86, p < 2.2e−16
R = −0.63, p < 2.2e−16
R = 0.81, p < 2.2e−16
R = 0.11, p = 0.04
R = 0.074, p = 0.16
R = −0.6, p < 2.2e−16
R = −0.31, p = 7.5e−10
R = 0.58, p < 2.2e−16
R = 0.76, p < 2.2e−16
R = −0.56, p < 2.2e−16
Average GC ratio (/bin)
Average CAI value (/bin)
Average AMFE (/bin)
Average half-life (/bin)
Average Length (nt) (/bin)

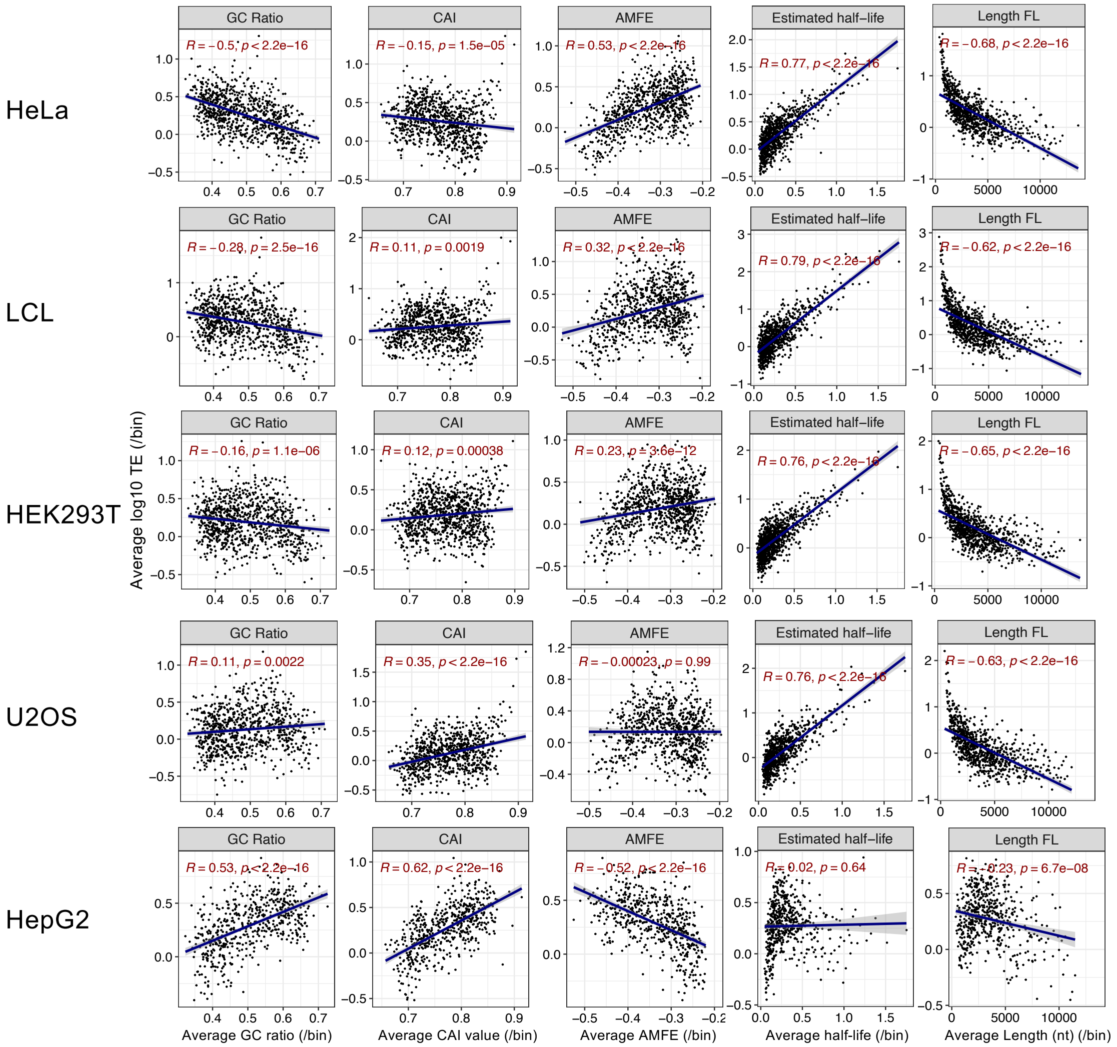


**Supplementary Figure 9. Correlation between sequence features and experimentally measured TE across multiple cell types.**

(**A**) Barplots showing the correlation between feature versus TE across 9 cell types. The y-axis shows the spearman's correlation coefficients (ρ) and the significance levels are labelled beside the bars.

(**B**) Scatter plots showing relationships between mRNA GC ratio, CAI, AMFE, estimated half-life, and mRNA length versus experimentally measured TE in HMECs, T47D, PC3, PC9, HeLa, LCL, HEK293T, U2OS, and HepG2. Data were binned by mRNA (10 mRNAs per bin), with each point representing the bin-averaged feature value and bin-averaged TE. Pearson's coefficients (R) and p-values are indicated.

**Supplementary Tables**

Supplementary Table 1. TE-altering gnomAD variants.

Supplementary Table 2. RNA structural changes of TE-altering ExAC variants.

Supplementary Table 3. RBP binding score changes of TE-altering ExAC variants.

Supplementary Table 4. TE-altering GWAS variants.

Supplementary Table 5. TE-altering ClinVar variants.

Supplementary Table 6. TE-altering ExAC variants shared in more than 2 cell lines.

Supplementary Table 7. TE-altering GWAS variants across cell lines.
